# Evolutionary safety of death by mutagenesis

**DOI:** 10.1101/2022.03.10.483790

**Authors:** Gabriela Lobinska, Yitzhak Pilpel, Martin A Nowak

**Affiliations:** Department of Molecular Genetics, Weizmann Institute of Science, Rehovot 76100, Israel; Department of Mathematics, Department of Organismic and Evolutionary Biology, Harvard University, Cambridge MA 02138, USA

## Abstract

Nucleoside analogs are a major class of antiviral drugs. Some act by increasing the viral mutation rate causing “death by mutagenesis” of the virus. Their mutagenic capacity, however, may lead to an evolutionary safety concern. We define evolutionary safety as a probabilistic assurance that the treatment will not generate an increased number of epidemiologically concerning mutated virus progeny. We develop a mathematical framework to estimate the total mutant load produced with and without mutagenic treatment. We predict rates of appearance of virus mutants as a function of the timing of treatment and the immune competence of patients, employing various assumptions about the vulnerability of the viral genome and its potential to generate undesired phenotypes. We focus on the case study of Molnupiravir, which is an FDA-approved treatment against COVID-19. We estimate that Molnupiravir is narrowly evolutionarily safe, subject to the current estimate of parameters. Evolutionary safety can be improved by restricting treatment to individuals with a low clearance rate and by designing treatments that lead to a greater increase in mutation rate. We report a simple rule to determine the fold-increase in mutation rate required to obtain evolutionary safety which is also applicable to other pathogen-treatment combinations.

## Introduction

Nucleoside analogs are molecules similar in shape to naturally occurring nucleosides used by living organisms and viruses for nucleic acid synthesis. They are therefore readily incorporated into nascent DNA or RNA chains by viral polymerases. Many nucleoside analogs differ from natural nucleosides in key aspects which usually prevents further viral genome chain elongation. Such nucleoside analogues lack a 3’OH group which makes the viral polymerase unable to attach the next nucleoside to the growing chain. Others, such as lamivudine, are D-enantiomers of natural nucleosides, and cause steric hindrance upon incorporation into the DNA or RNA chain^1^.

Other nucleoside analogues do not prevent viral transcription. Instead, they have the capacity to ambiguously base pair with several nucleosides. Therefore, they cause erroneous incorporation of nucleosides during the transcription process, thereby increasing the virus mutation rate up to the point of “death by mutagenesis”, a mechanism with foundations in quasispecies theory. This theory describes populations of replicating genomes under mutation and selection^2–7^.

Molnupiravir, which has been suggested as treatment against COVID-19, seems to act exclusively through mutagenesis. Its incorporation into nascent RNA genomes by the viral polymerase does not result in chain termination: in fact, the viral RNA polymerase has been shown to successfully elongate RNA chains after the incorporation of Molnupiravir^8–10^. Molnupiravir switches between two tautomeric forms: one is structurally similar to a cytosine, the other is structurally similar to a uracil. Hence, Molnupiravir can base pair, depending on its form, either with guanosine or with adenosine^8,9^. SARS-COV2 is a positive-sense single-stranded RNA virus and its RNA replication proceeds in two steps. First, the negative-sense RNA is polymerized based on the plus strand, and the negative strand then serves as a template to synthetize positive-sense RNA molecules^11^. Hence, the incorporation of Molnupiravir during the first step of RNA synthesis gives rise to an ambiguous template: positions where Molnupiravir was incorporated can be read by the RNA-dependent RNA polymerase as either guanosine or adenosine. This causes mutations in the progeny RNA compared with the parental RNA, possibly up to the point of the “error catastrophe” and death of the virus^8–10^.

While considerable theoretical basis describes death by mutagenesis, a theoretical treatment is still missing to describe quantitatively the potential of emergence of variants of concern (VoC) upon mutagenesis. For example, in the context of the COVID-19 pandemic, the mutagenic potential of Molnupiravir naturally causes concerns about accelerating SARS-COV2 evolution. The evolution of resistance of SARS-COV2 against vaccination or existing treatments as well as enhanced transmissibility or lethality is a major concern, which has given rise to an impressive number of studies^12-27^ that have been founded on a long history of modelling the emergence of resistance against treatment in other viruses^28–36^ and epidemiological models for disease spread^18,19,37,38^.

As noted before, the intended antiviral activity of Molnupiravir resides in its capacity to induce mutagenesis and hence reduce virus load. Yet, this very property which confers to Molnupiravir its desired antiviral effect might also enhance the capacity of the virus to develop drug resistance, immune evasion, infectivity or other undesired phenotypes. Thus, a mathematical analysis should weigh the desired and potentially deleterious effects of mutagenesis drugs in general, and of the present virus and drug in particular.

In this paper, we analyze the case study of the increase of the evolutionary potential of a virus (here: SARS-COV2) under mutagenic treatment (here: Molnupiravir treatment). In particular, we ask if the wanted effect of limitation of virus load by the drug could be accompanied by an unwanted enhancement in the rate of appearance of new VOCs due to increased mutagenesis. We construct a mathematical framework describing the increase and decrease of the virus load after infection and derived expressions for the total amount of wild-type and mutant produced by individuals during the course of an infection. We use empirical data on COVID-19 and bioinformatic data on SARS-COV2 to estimate key parameters, including infection progression within the body amidst response of the immune system and the number of potentially lethal positions in the genome.

We find that the Molnupiravir-SARS-COV2 couple is situated in a region of the parameter space which is narrowly evolutionarily safe. Evolutionary safety increases with decreasing clearance rate in treated patients and with higher number of viral genome positions that are lethal when mutated. Crucially, evolutionary safety could be improved by obtaining higher increases in the mutation rate under treatment which provides a clear direction for drug improvement. We suggest a simple mathematical formula which determines the evolutionary safety of a drug given the pathogen’s mutation rate with and without treatment and the number of positions in the pathogen’s genome that are lethal when mutated.

### Description of the model

After infection with SARS-COV2, virus load increases exponentially until it reaches a peak after a median of about 5 days^39^. During this growth phase the action of the immune system is insufficient to counterbalance viral replication. Subsequently the immune response gains momentum and infection enters a clearance phase. Now virus load decreases exponentially until the virus becomes eliminated about 10-30 days after initial infection^35,39^. In some immunocompromised individuals, viral clearance can take many weeks^40,41^. However, some argue that the isolation of infectious virus is rare after 20 days post-infection^42^.

In our mathematical formalism, we describe the evolution of a virus within the body of a single human host by following the abundance of two viral types: wild type, *x*, and concerning mutants, *y*. Concerning mutants are those that can lead to undesirable viral evolution, for example to escape from vaccination^14^ or toward higher virulence or infectivity. Later we broaden the definition of variable *y* to include any viable mutant, as these may subsequently facilitate epistatic tracks towards to VOCs. Both *x* and *y* replicate with birth rate *b* and replication quality *q* = 1 − *u*, where *u* is the mutation rate per base. The mutation rate can be altered by the administration of a mutagenic drug. The virus genome contains *m* positions, all of which must be maintained without mutations in order to generate viable progeny. We consider *n* positions, such that even a single mutation in one of them gives rise to a concerning mutant virion, *y*. As common in mutagenesis and also in the specific mechanism of action of Molnupiravir, transition mutations are more likely than transversion mutations (see **Figure 1A**). Our model can be extended to consider situations where the mutagenic drug increases the probability of mutation for a subset of all possible mutations (see **Methods**). Both *x* and *y* are cleared at same rate *a*_*j*_ with the subscript *j* indicating the presence or absence of an adaptive immune response. During the growth phase *j* = 0 and during the clearance phase *j* = 1. We have *a*_0_ < *b* < *a*_1_. Virus dynamics ^43^ in an infected patient can be described by the system of differential equations

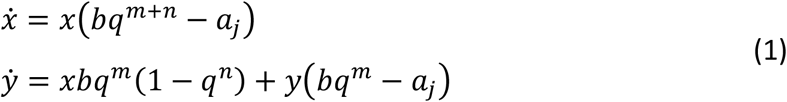

**Figure 1.**
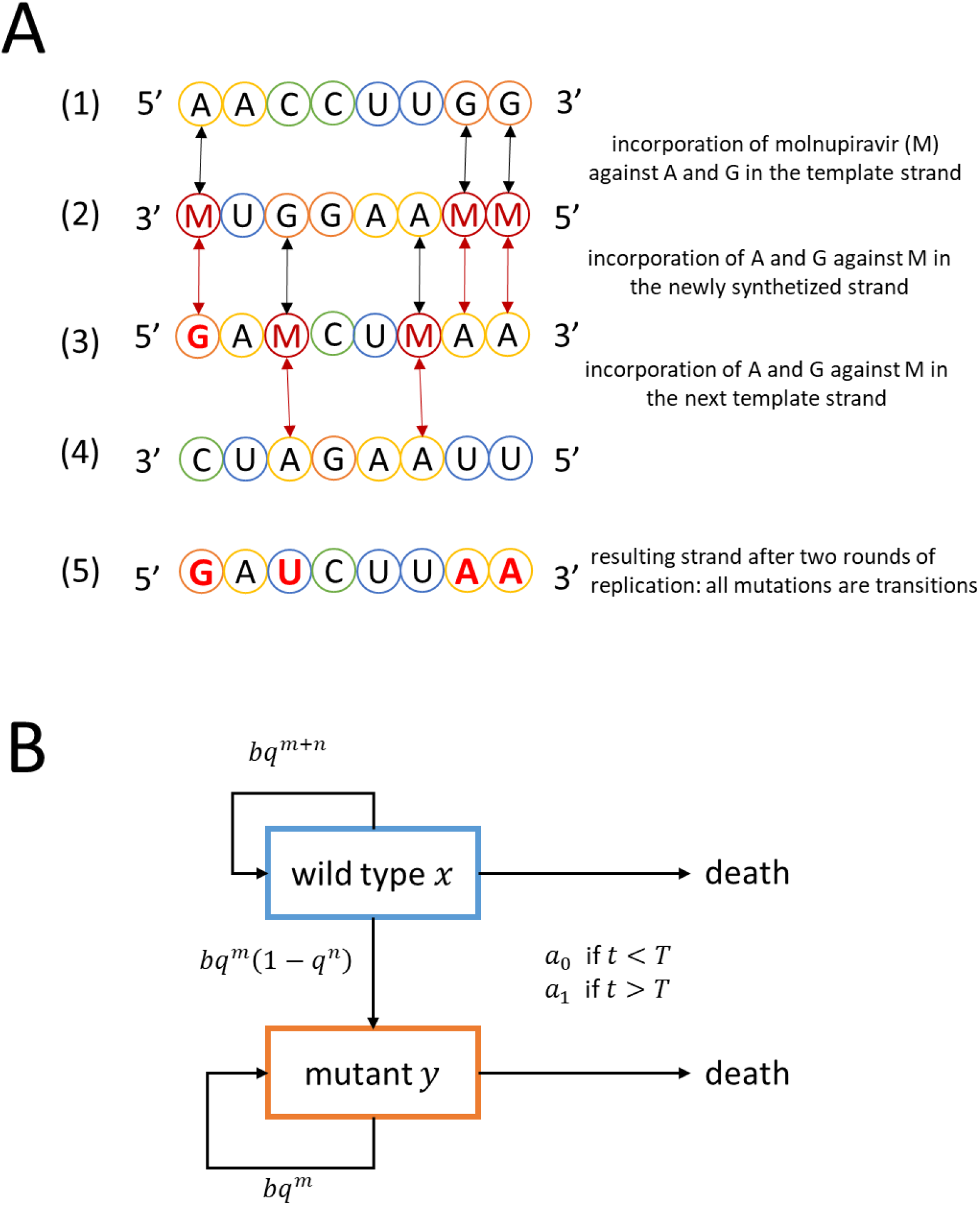
A: Mechanism of action of molnupiravir. SARS-COV2 has a positive-sense single-stranded RNA genome, represented schematically in (1). Its replication proceeds by two steps: first, the synthesis of a negative-sense template strand (2), which is then used to synthesize a positive-sense progeny genome (3). Molnupiravir (designated by M) is incorporated against A or G during the synthesis of the negative-sense template strand (2). When the template strand is replicated, M can be base-paired with either G or A. Hence, all A and G in the parent genome become ambiguous and can appear as A or G in the newly synthetized positive-strand genome; see position 1 in (3). C and T are not affected by molnupiravir during the synthesis of the template strand, (1) to (2), but can be substituted to M during the synthesis of the progeny genome from the template strand; see (2) to (3). As previously, M can then base-pair with A or G when used as a template; see (3) to (4), which can cause A->U and U->A transitions in the final progeny genome (5). **B: Virus dynamics within an infected person**. Wildtype (*x*) and the mutant (*y*) replicate at rate *b* and quality *q* = 1 − *u*. The per base mutation rate, *u*, is increased by treatment with molnupiravir. Both the wildtype and the mutant need to maintain *m* positions to remain viable. Mutating any of *n* positions in the wildtype results in a mutant. In the beginning of the infection, the adaptive immune response is weak, and the virus is cleared at a rate *a*_0_ which is less than *b*. After some time, *T*, the adaptive immunity is strong, and the virus is cleared at the higher rate *a*_1_ which is greater than *b*.

We ignore back mutation from mutant to wild type^2,5,43^. In the growth phase, without treatment, we have *bq*^*m*+*n*^ > *a*_0_ since both *x* and *y* grow exponentially. In the clearance phase, without treatment, we have *bq*^*m*^ < *a*_1_ since both *x* and *y* decline exponentially. The system is linear and can be solved analytically (see **Methods**). The biological reactions are presented schematically in **Figure 1B**. In our simple approach, there is a sudden onset of adaptive immunity which happens at time *T*. We relax this assumption in a model extension.

### Estimating parameters

#### Estimation of mutation rates

All parameters and methods for their estimation are summarized in **Table 1**. Each parameter can be estimated from existing literature. We denote by *u*_0_ the mutation rate without mutagenic treatment and by *u*_1_, which is greater than *u*_0_, the mutation rate with mutagenic treatment.

**Table 1.** Summary of parameters with ranges for their values and method of estimation.

| Symbol | Name | Value | Method of approximation | References |
| --- | --- | --- | --- | --- |
| $b$ | birth rate of infected cells | 7.61 | fitted to virus load along time measurements in infected patients | 39 |
| $a_0$ | clearance rate prior to adaptive immune response | 3 | computed from eclipse time of SARS-COV2 in infected cells *in vitro* | 47 |
| $a_1$ | clearance rate during adaptive immune response | 7.7 – 10 | fitted to virus load along time measurements in infected patients | 35,39–42 |
| $u_0$ | viral mutation rate without treatment | $10^{-6}$ | mutation rate measured for related MHV | 47,48 |
| $u_1$ | viral mutation rate during treatment | $2 - 5 \cdot 10^{-6}$ | fold-increase in mutation rate under treatment measured in treated patients and *in vitro* | 10 |
| $m$ | number of lethal positions in SARS-COV2 genome | ~12,000 | typical proportion of lethal mutations in ssRNA viruses | 46 |
|  |  | ~21,000 | typical proportion of lethal + severely deleterious mutations in ssRNA viruses |  |
| $n$ | number of beneficial positions in SARS-COV2 genome | ~100 | analysis of mutagenesis data | 54,55 |
| $T$ | time of peak of virus load | 3-7 | virus load along time measurements in infected patients | 39 |

The typical mutation rate for other positive single-strand RNA viruses is 10^−544^. The mutation rate of SARS-COV2 has been hypothesized to be lower because of a proofreading capability^45^. The per-base mutation rate has been estimated at *u*_0_ = 10^−6^ by proxy with the related beta-coronavirus MHV^46,47^. An in vitro study of experimental evolution of SARS-COV2 has reached the estimate *u*_0_ = 3.7 · 10^−648^. Another study measuring the mutation rate of SARS-COV2 *in vitro* has estimated *u*_0_ = 2.5 · 10^−510^. For our analysis, we use *u*_0_ = 10^−6^.

The mutation rate of SARS-COV2 under Molnupiravir treatment has been measured *in vitro* to be 2 to 5-fold higher than without treatment^10^. The fold-increase in mutation rate under treatment can also be estimated from sequencing viral samples from treated patients. A 2-fold increase in the mutation rate in RNA-dependent RNA polymerase sequence in patients treated with Molnupiravir has been observed during its phase 2a clinical trial^49^. This estimate comes with the caveat of neglecting potentially rare, severely deleterious mutants since those are less likely to be sequenced. Hence, we estimate *u*_1_ to be 2 to 5 times higher than *u*_0_. Mutation rate estimations for different pathogen-drug combinations are available in the literature, and result in even higher estimates for the virus mutation rate under treatment^50^. In our analysis we explore a wide range of *u*_1_ values, because it is our expectation that future mutagenic treatments might achieve higher increases of the virus mutation rate.

#### Estimations of viral birth and clearance rates

The average lifetime of an infected cell is about 8 hours^47^. Hence, without infection of new cells we would obtain a clearance rate of *a*_0_ = 3 per day. From the current literature, we know that the virus load grows by about 10 orders of magnitude within 5 days^39,51^. Hence, for the viral growth rate we obtain *b* = 7.61. For the clearance phase, a decrease by 4 orders of magnitude in 10 days results in a death rate of *a*_1_ = 8.76 per day reflecting high immunocompetence. The same decrease over 120 days results in a death rate of *a*_1_ = 7.69 per day reflecting low immunocompetence (see **Methods**). These estimates are approximations as they ignore loss by lethal mutants.

#### Estimation of the number of viral genome positions that are either lethal or potentially concerning when mutated

The distribution of fitness effects of random, single mutations has been studied in a different single-stranded RNA virus, the vesicular stomatitis virus (VSV)^52^. This distribution seems to be similar among single-stranded RNA viruses but could differ between species^53^. According to these studies, the proportion of viral genome positions that are lethal when mutated is about 40% and the proportion of highly deleterious mutations, defined as those that reduce the viral fitness by more than 25%, represents about 30%. Note that the small mutation rate allows us to approximate the number of lethal positions as 1/3 of the total number of possible mutations, taking into account that each position can be mutated to three different destinations. SARS-COV2 genome has a length of 29,900 nt. Hence, we have *m* = 11,960 when considering lethal mutations only and *m* = 20,930 when considering both lethal and highly deleterious mutations. Hence, the realistic range for *m* is between 11,960 and 20,930. For completeness, we also explore unrealistically low values of *m* such as 1,500, which is the number of positions in the coding genome that are one nucleotide way from a STOP codon.

In order to estimate the number of positions that could give rise to new variants of concern when mutated (denoted by *n*), we used empirical data collected by^54,55^. Starr et al. conducted deep mutagenesis scans of the receptor-binding domain of the SARS-COV2 spike protein. For each of the generated mutants, Starr et al. measured the mutant’s binding affinity to ACE2 which is the receptor used by SARS-COV2 to enter the human cell. In a subsequent study, Starr et al. also measured each mutant’s affinity to antibodies in order to assess the ability of each mutant to escape the adaptive immune response and antibody treatments. Both escape from antibody and increased affinity to ACE2 are phenotypes beneficial for SARS-COV2. We identified 484 amino acid substitutions that result in antibody escape and 314 distinct amino acid substitutions that result in increased binding to ACE2 (see detailed information in the **Methods**). For each position coding for the receptor-binding domain of the spike protein, we counted how many mutations can give rise to the identified set of beneficial substitutions (we corrected for the overlap of substitutions found in both categories). We found that the resulting estimate (divided by 3 to take into account all possible destinations, see **Methods**) was *n* = 87 when considering all possible mutations and *n* = 75 when considering only transition mutations, i.e. when taking into account the specific mechanism of action of Molnupiravir.

Of course, mutations that are advantageous for the virus could occur also outside of the receptor-binding domain of the spike protein. More broadly, any neutral and even slightly deleterious mutation can be undesirable since they could represent an evolutionary “stepping-stone” to a multiple-mutation variant due to epistasis. Hence, we also explore how considering a very large number of positions that could give rise to new variants of concern when mutated, up to the length of the SARS-COV2 genome minus the *m* positions that are lethal when mutated.

### Abundance of mutant virus for various treatment regimes

In **Figure 2**, we show the dynamics of total virus and mutant over the course of an infection. We consider four times for the start of mutagenic treatment: at infection; at day 2 after infection, which corresponds to the beginning of symptoms; at day 5 after infection, which corresponds to the peak of the virus load; and at day 7 after infection. We observe that treatment always decreases the abundance of wild type virus. The dynamics of mutant follows that of the wild type. For the parameters used in **Figure 2**, treatment decreases the abundance of mutant virus – with exception of a brief transient period soon after the start of therapy, which is almost invisible in the figure.

**Figure 2.**
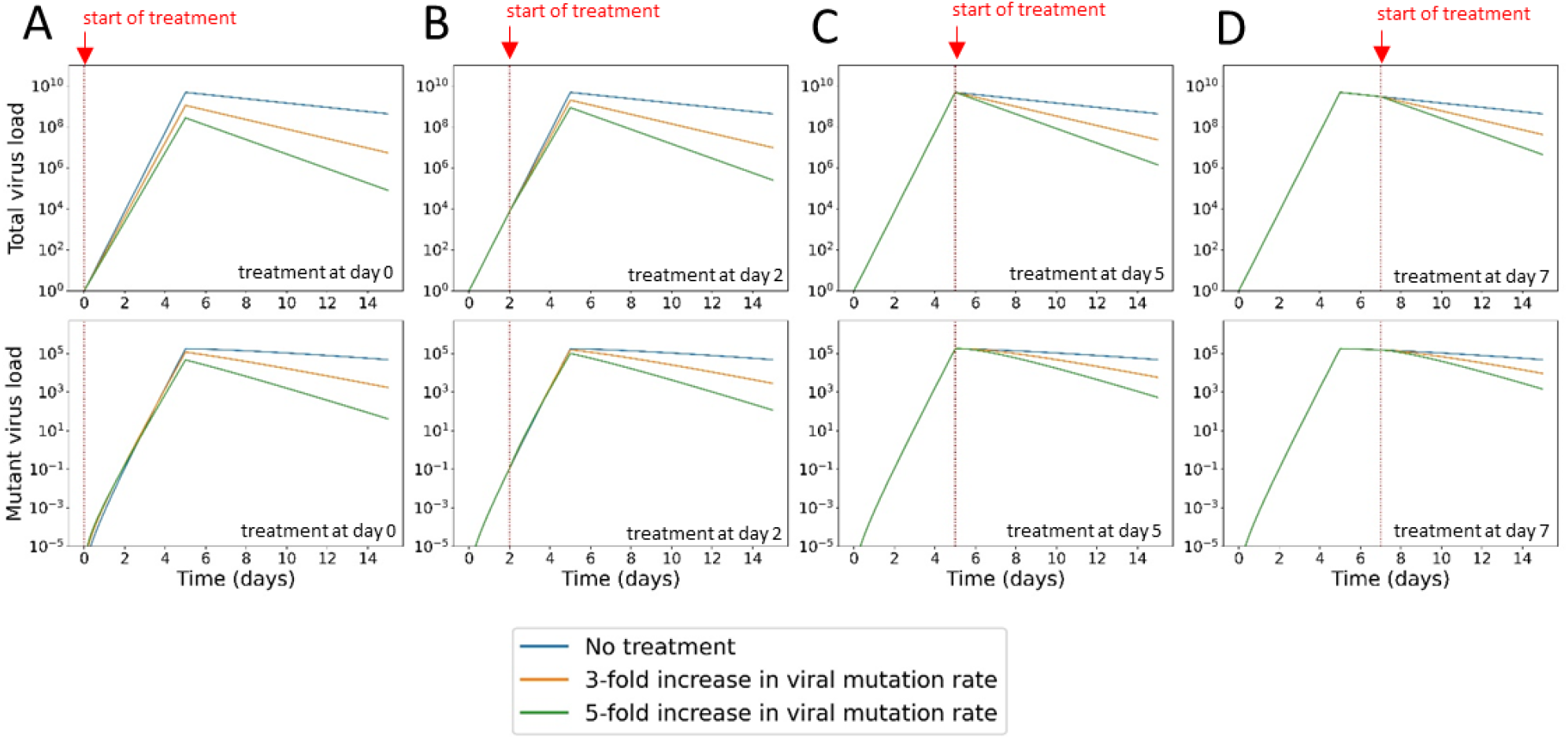
Time series of virus (*v* = *x* + *y*) and mutant (*y*) virus with and without treatment. We consider four starting points for treatment: (A) at infection, at day 0; (B) before reaching peak virus load, at day 2; (C) at peak virus load, day 5; and (D) after peak virus load, day 7. The red arrow and dotted line indicate the beginning of treatment. Virus load increases during the first 5 days when the death rate is *a*_0_. Virus load subsequently decline when the death rate is *a*_1_. We observe that treatment with a mutagenic drug reduces the total abundance of virus. The higher the mutation rate induced by treatment, the higher is the decrease in virus load. The abundance of mutant virus can increase transiently after the start of treatment, but subsequently declines compared to the case of no treatment. Parameters: *a*_0_ = 3, *a*_1_ = 7.7, *b* = 7.61, *u*_0_ = 10^−6^, *m* = 20,000, *n* = 1. Initial condition: *x*_0_ = 1 and *y*_0_ = 0.

We are now interested in calculating the total number of mutant virus produced over the course of infection. This number can be computed as the integral of the abundance of mutant virus over time (see **Methods**). We consider two scenarios: in the first, the patient begins treatment when their virus load reaches its peak; in the second, the patient begins treatment when they become infected (following exposure to an infected individual).

#### Treatment begins at (or near) peak virus load

In **Figure 3**, we show the cumulative mutant load, *Y*(*u*_1_), as a function of the mutation rate *u*_1_ for the case where treatment starts at peak virus load. To understand this function, we introduce the parameter *η* = *x*_*T*_/*y*_*T*_, with *x*_*T*_ and *y*_*T*_ denoting respectively wildtype and mutant virus load at peak, which is reached for each strain at a time. If *η* > *n*/*m* then *Y*(*u*_1_) is a declining function. In this case, any mutagenic treatment is evolutionarily safe in the sense of reducing the cumulative mutant virus load. If on the other hand *η* < *n*/*m* then the function *Y*(*u*_1_) attains a single maximum at

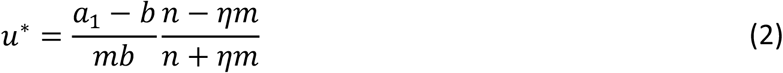

**Figure 3.**
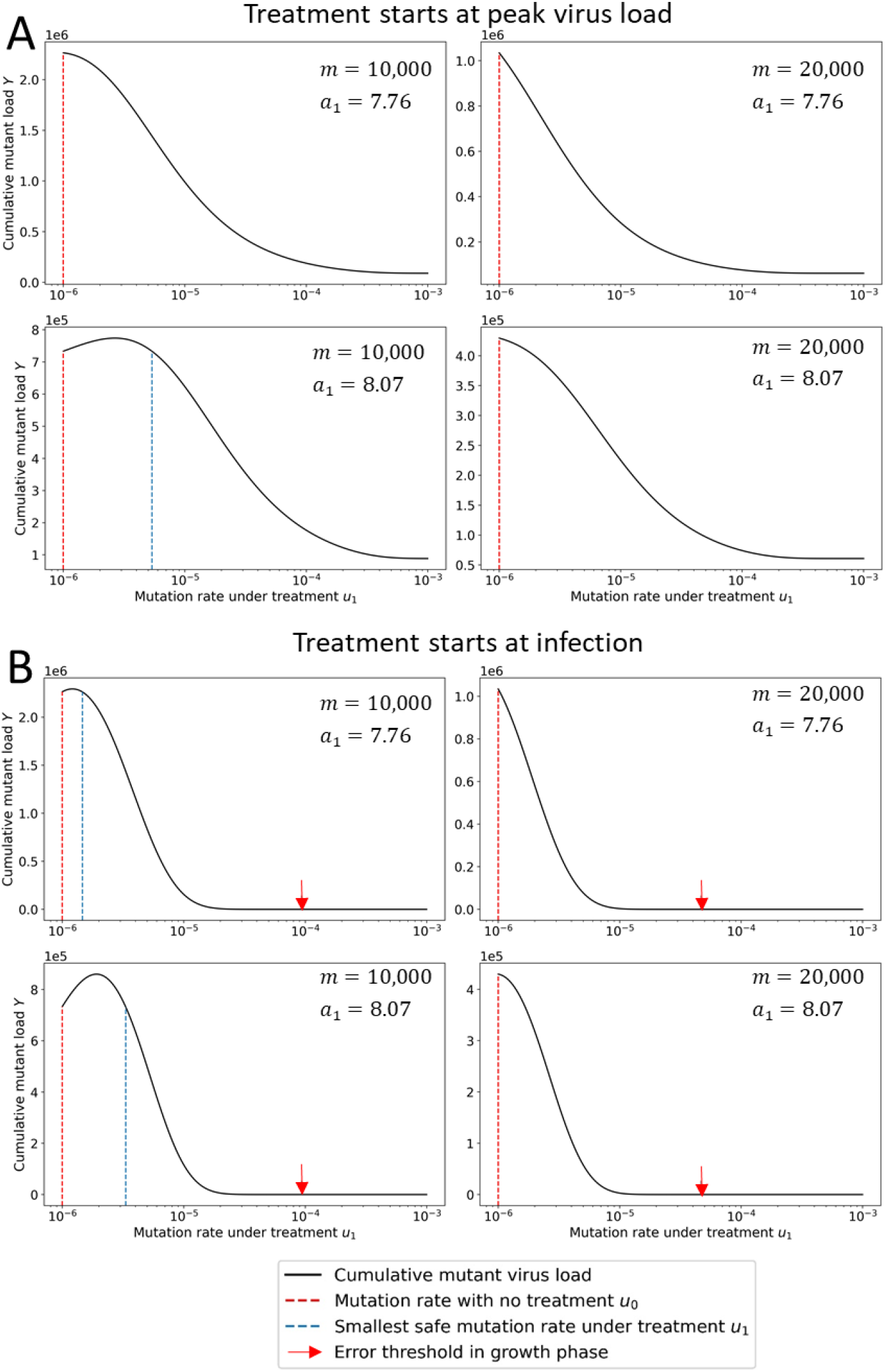
Cumulative mutant virus load versus mutation rate, *u*_1_, during treatment. The cumulative mutant virus load increases with mutation rate *u*_1_ before reaching a peak and then decreases to low values. If the peak is reached at a mutation rate that is less than the natural mutation rate, *u*_0_ (red dotted line), then any increase in mutation rate reduces the cumulative mutant load. If the peak is reached for a mutation rate greater than *u*_0_, then the increase in mutation rate caused by mutagenic treatment must exceed a threshold value (blue dotted line) to reduce the cumulative mutant virus load. (A) Treatment starts at peak virus load. (B) Treatment starts at infection. The red arrow indicates the mutation rate at the error threshold of the growth phase. Parameters: *b* = 7.61, *a*_0_ = 3, *n* = 1, *T* = 5, *m* and *a*_1_ as shown.

If *u*_0_ > *u*^*^ then any increase in mutation rate is beneficial as it actually *decreases* the chance of appearance of VOC compared to evolution of the virus under no treatment. If *u*_0_ < *u*^*^ then a small increase in the mutation rate can increase the chance of appearance of VOC under treatment, and thus be evolutionarily unsafe; in this case there needs to be a sufficiently large increase in mutation rate to make the treatment evolutionarily safe (see **Figure 3** for details). We notice that increasing estimates of *m* or decreasing *a*_1_ reduces the value of *u*^*^ and therefore increases the range of *u*_0_ for which mutagenic treatment is evolutionarily safe. In particular, the more immunocompromised a patient is (lower *a*_1_), the lower the value of *u*^*^. In **Figure 3**, we notice that only for low *m* and high *a*_1_ we find *u*^*^ > *u*_0_. For all other cases, *u*^*^ > *u*_0_, and death by mutagenesis is both evolutionarily safe and evolutionarily desired, because it reduces the abundance of both wild type and mutant.

#### Treatment begins at (or soon after) infection

In **Figure 3**, we also show the cumulative mutant load, *Y*(*u*_1_), as a function of the mutation rate *u*_1_ for the case where treatment starts at infection. We find that this function attains a maximum at a value which is given by the root of a third order polynomial (see **Methods** and **Extended Data Figure 1**). Using the notation *k* = [*b*(2*b* − *a*_0_ − *a*_1_)]/[(*b* − *a*_0_)(*a*_1_ − *b*)] and *h* = *bT*, we can approximate *u*^*^ as follows:

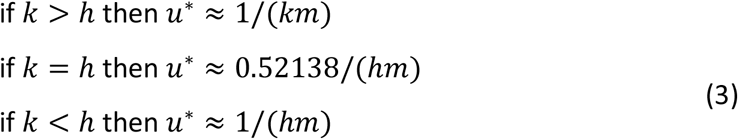

Again if *u*_0_ > *u*^*^ then any increase in mutation rate is beneficial. If *u*_0_ < *u*^*^ then a small increase in the mutation rate can be evolutionarily not safe, but a sufficiently large increase in mutation rate can make the treatment evolutionarily safe (see **Figure 3** for more details).

#### Exploring the parameter space for evolutionary safety

In **Figure 4**, we show the fold-increase in virus mutation rate that mutagenic treatment has to achieve to be evolutionarily safe. We vary first the number of lethal mutations *m* in the viral genome and the clearance rate *a*_1_. For treatment starting at peak virus load (**Figure 4A**), we find that increase in mutation rate is evolutionarily safe if *m* > 22,000 or *a*_1_ < 7.8 (green region). Evolutionary safety becomes an issue for small values of *m* and larger values of *a*_1_. For *m* = 12,000 and *a*_1_ = 9 we need at least a 10-fold increase in mutation rate before the drug attains evolutionary safety. When treatment begins at infection (**Figure 4B**) the evolutionarily safe area becomes smaller, but the minimum increase in mutation rate required for evolutionary safety is lower. For example, for *a*_1_ = 9 and *m* = 12,000, we need only a 3-fold increase. We show the same figure, but for an extended range of *m* values in **Extended Data Figure 2**.

**Figure 4.**
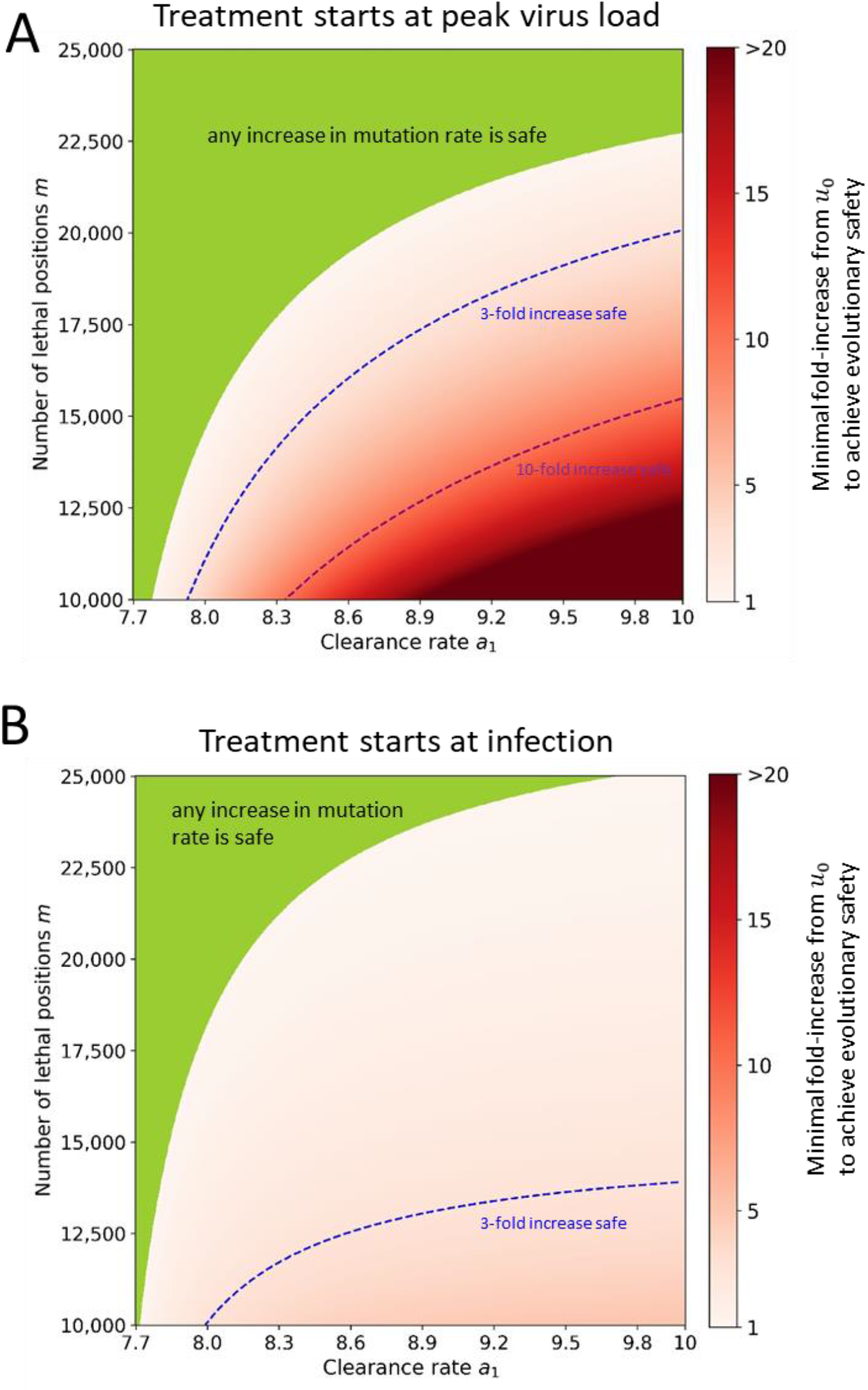
Evolutionary safety of mutagenic treatment. In the green parameter region, any increase in mutation rate reduces the cumulative mutant virus and is therefore evolutionarily safe. In the red shaded region, we indicate the minimum fold increase in mutation rate that is required to reduce the cumulative mutant load. Contour lines for 3-fold and 10-fold increase are shown. (A) Treatment starts at peak virus load. (B) Treatment starts at infection. Parameters: *b* = 7.61, *a*_0_ = 3, *n* = 1, *T* = 5, *u*_0_ = 10^−6^.

#### Evolutionary risk factor (ERF) and infectivity risk factor (IRF)

We define the “evolutionary risk factor” (ERF) of mutagenic treatment as the ratio of cumulative mutant virus load with treatment compared to without treatment (see **Methods**). The condition for evolutionary safety of mutagenic treatment is that ERF is less than one. Denote by *Y*_*ij*_ the cumulative mutant load with the subscript *i* indicating the presence (*i* = 1) or absence (*i* = 0) of treatment during the growth phase, and the subscript *j* indicating the presence (*j* = 1) or absence (*j* = 0) of treatment during the clearance phase. Therefore, *Y*_00_ is the cumulative mutant load without treatment, *Y*_01_ is the cumulative mutant load with treatment in the clearance phase, and *Y*_11_ is the cumulative mutant load with treatment in both growth and clearance phase. For treatment that starts at peak, *ERF* = *Y*_01_/*Y*_00_. For treatment that starts at infection, *ERF* = *Y*_11_/*Y*_00_. An evolutionary risk factor below one signifies that treatment reduces the mutant load, and hence treatment can be even encouraged from an evolutionary perspective. An evolutionary risk factor above one implies that treatment increases the mutant load.

In addition, we define the “infectivity risk factor” (IRF) which quantifies the efficacy of the treatment. The IRF is the ratio of the total cumulative viral load, mainly governed by the wild-type, with treatment compared to the total cumulative viral load without treatment. IRF is always below 1.

In **Table 2**, we computed some values for the cumulative mutant load with and without treatment and the cumulative total virus load with and without treatment, as well as the corresponding ERF and IRF. We notice that ERF increases (hence evolutionary safety decreases) with clearance rate, *a*_1_. However, both the cumulative mutant viral load with and without treatment decrease with clearance rate. Hence, although the ERF is higher for more immunocompetent individuals, the absolute quantity of mutant produced is lower. We also notice that the IRF increases with immunocompetence, indicating that the benefit of treatment is smaller for more immunocompetent individuals who clear the virus rapidly.

**Table 2.**
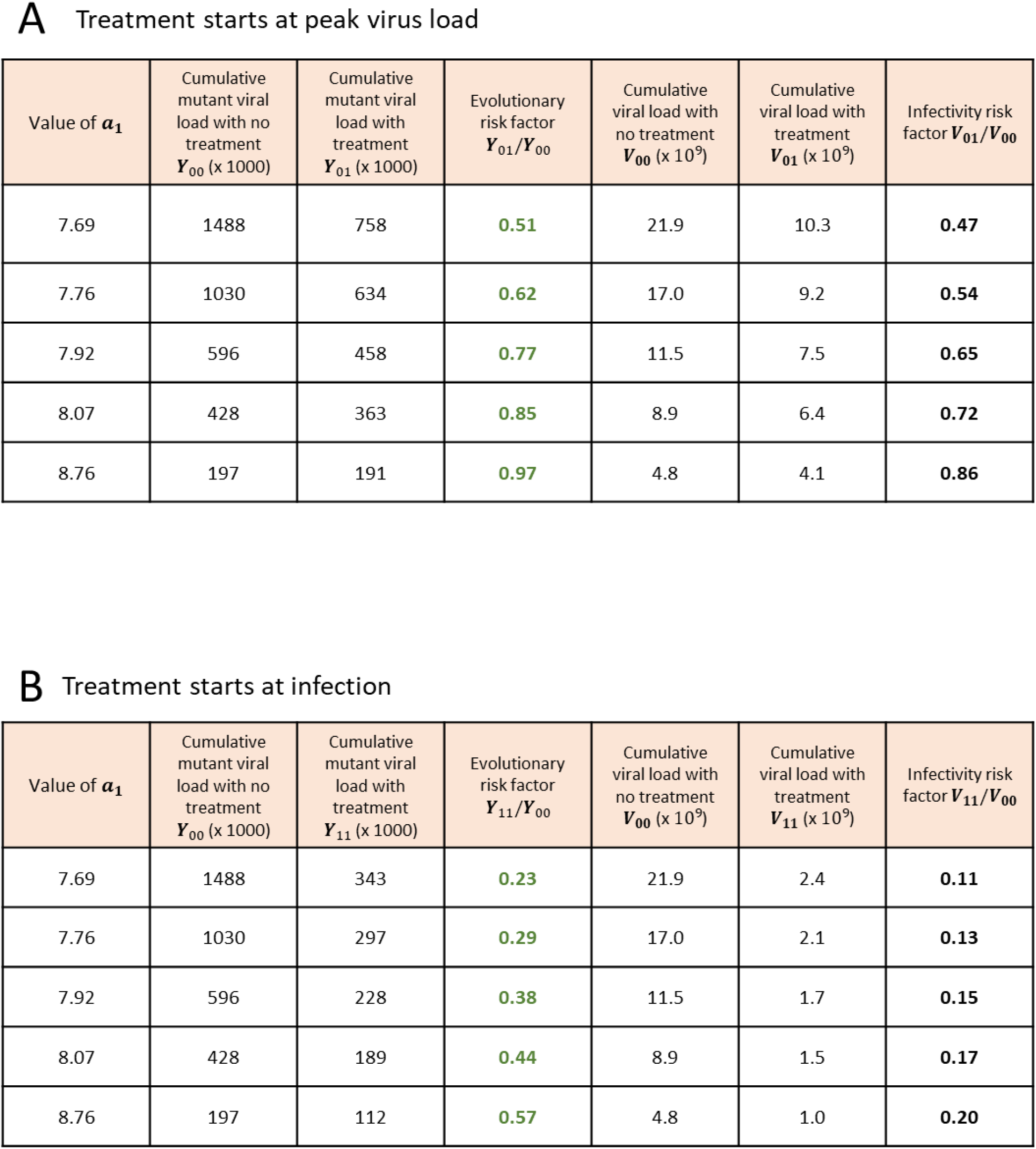
Cumulative virus load, mutant load, infectivity risk factor (IRF) and evolutionary risk factor (ERF) of mutagenic treatment. (A) Treatment starts at peak virus load. (B) Treatment starts at infection. We show numerical results for individuals that differ in their immune competence, which affects the clearance rate, *a*_1_, during adaptive immunity. Patients that are less immunocompetent benefit more from mutagenic treatment (lower IRF) and also have a lower ERF. Parameters: *a*_0_ = 3, *b* = 7.61, *u*_0_ = 10^−6^, *u*_1_ = 3 · 10^−6^, *m* = 20,000, *n* = 1, *T* = 5. Initial condition: *x*_0_ = 1 and *y*_0_ = 0.

A Treatment starts at peak virus load
| Value of $a_1$ | Cumulative mutant viral load with no treatment $Y_{00}$ (x 1000) | Cumulative mutant viral load with treatment $Y_{01}$ (x 1000) | Evolutionary risk factor $Y_{01}/Y_{00}$ | Cumulative viral load with no treatment $V_{00}$ (x $10^9$) | Cumulative viral load with treatment $V_{01}$ (x $10^9$) | Infectivity risk factor $V_{01}/V_{00}$ |
| --- | --- | --- | --- | --- | --- | --- |
| 7.69 | 1488 | 758 | 0.51 | 21.9 | 10.3 | 0.47 |
| 7.76 | 1030 | 634 | 0.62 | 17.0 | 9.2 | 0.54 |
| 7.92 | 596 | 458 | 0.77 | 11.5 | 7.5 | 0.65 |
| 8.07 | 428 | 363 | 0.85 | 8.9 | 6.4 | 0.72 |
| 8.76 | 197 | 191 | 0.97 | 4.8 | 4.1 | 0.86 |

**Table 2. Cumulative virus load, mutant load, infectivity risk factor (IRF) and evolutionary risk factor (ERF) of mutagenic treatment.**
| Value of $a_1$ | Cumulative mutant viral load with no treatment $Y_{00}$ (x 1000) | Cumulative mutant viral load with treatment $Y_{11}$ (x 1000) | Evolutionary risk factor $Y_{11}/Y_{00}$ | Cumulative viral load with no treatment $V_{00}$ (x $10^9$) | Cumulative viral load with treatment $V_{11}$ (x $10^9$) | Infectivity risk factor $V_{11}/V_{00}$ |
| --- | --- | --- | --- | --- | --- | --- |
| 7.69 | 1488 | 343 | 0.23 | 21.9 | 2.4 | 0.11 |
| 7.76 | 1030 | 297 | 0.29 | 17.0 | 2.1 | 0.13 |
| 7.92 | 596 | 228 | 0.38 | 11.5 | 1.7 | 0.15 |
| 8.07 | 428 | 189 | 0.44 | 8.9 | 1.5 | 0.17 |
| 8.76 | 197 | 112 | 0.57 | 4.8 | 1.0 | 0.20 |

In **Figure 5** and **Extended Data Figure 3**, we explore the ERF for wider regions of the parameter space. We vary each pair of parameters, while fixing others at their most probable value. The ERF exceeds 1 when the number of positions that would be lethal when mutated is much lower than our minimum estimate (*m* < 12,000). As *m* decreases treatment induces less death by mutagenesis and thus provides more opportunity for mutants to be generated and to survive. Again, we observe that evolutionary safety decreases with the clearance rate, *a*_1_. Delaying treatment, especially past the peak of the virus load, brings ERF closer to 1. Hence, early treatment for high enough *m* should be encouraged since it can substantially decrease the abundance of mutant. Overall, we notice that most regions of the parameter space are evolutionarily safe.

**Figure 5.**
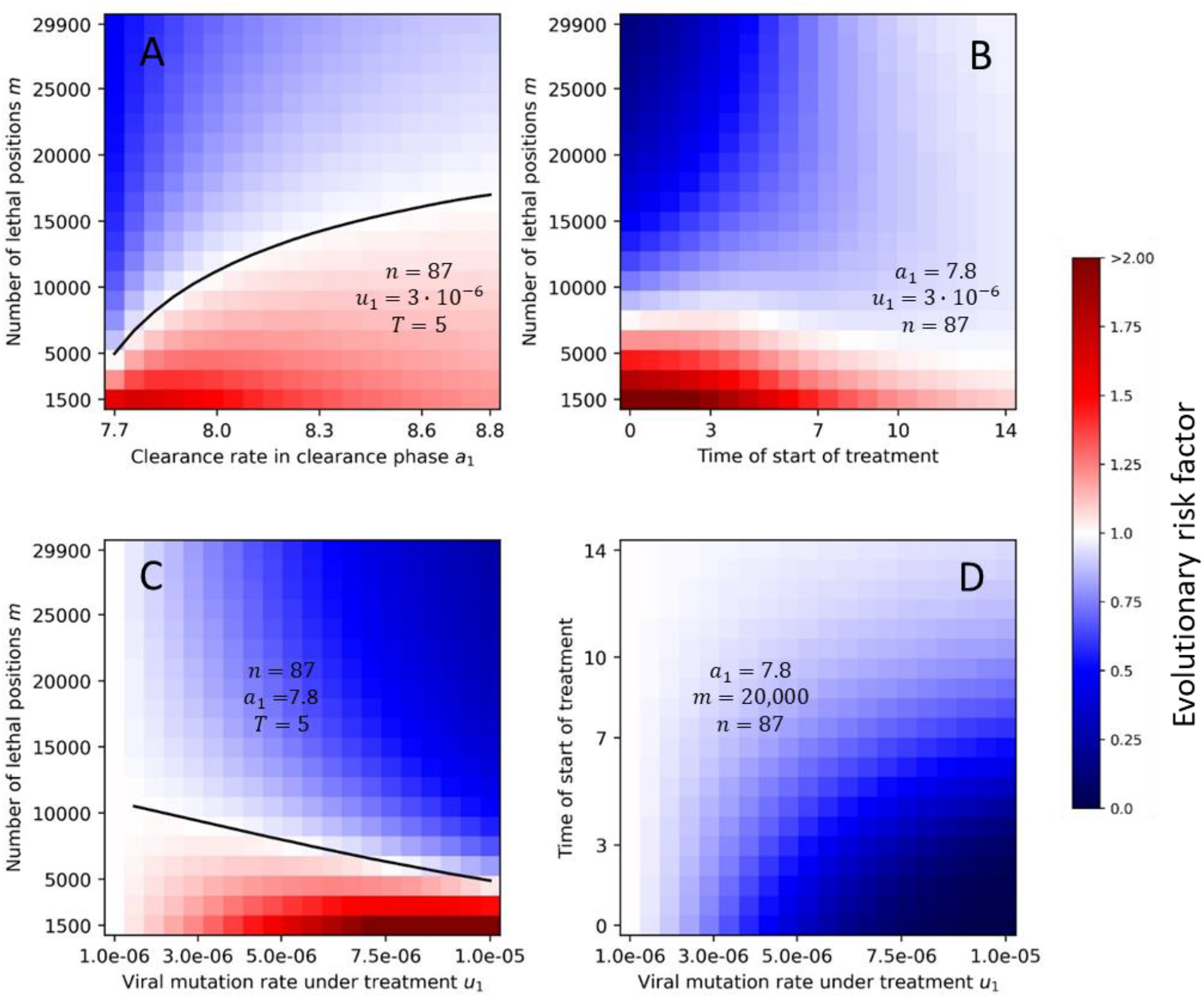
Evolutionary risk factor (ERF) for a grid of pairs of selected parameters. For each pair of parameters, we numerically compute the ERF for a range of values, while all other parameters are fixed. We observe that the value of *n* has little effect on the ERF. Evolutionary risk factors above 1 are only observed for low values of the number of lethal positions, *m*. The ERF decreases with early treatment, high viral mutation rate under treatment, and large number of lethal positions. Initial condition: *x*_0_ = 1 and *y*_0_ = 0.

In **Extended Data Figure 4**, we explore the ERF for lower and higher values of the birth rate *b* and the clearance rate *a*_0_ in the growth phase. We adjust the values of *b* and *a*_0_ such that the net growth rate is conserved (ignoring lethal mutations). We observe that smaller values of *b* and *a*_0_ lead to an increase in ERF, while larger values to a decrease.

### The evolutionary risk factor is a slowly declining function of the number of concerning mutations

So far, we have used the parameter *n* to denote the number of mutations which would result in variants of concern (VOCs) that is variants with increased transmissibility, virulence or resistance to existing vaccines and treatments. However, in the broad sense, any treatment which increases the standing genetic variation of the virus could favor the emergence of new variants of concern by enabling epistatic mutations. Therefore, we now extend the interpretation of *n* to include any viable mutation in the viral genome.

In **Figure 6**, we show that the ERF is a declining function of *n*. Thus, the more opportunities the virus has for concerning mutations (the larger *n*), the higher the advantage of mutagenic treatment. The reason for this counter-intuitive observation is that for large *n* the cumulative mutant virus load is high already in the absence of treatment, while mutagenic treatment reduces the mutant load by forcing additional lethal mutations. ERF decreases with the number of positions *n* also for lower birth rate *b* (**Extended Data Figure 5**).

**Figure 6.**
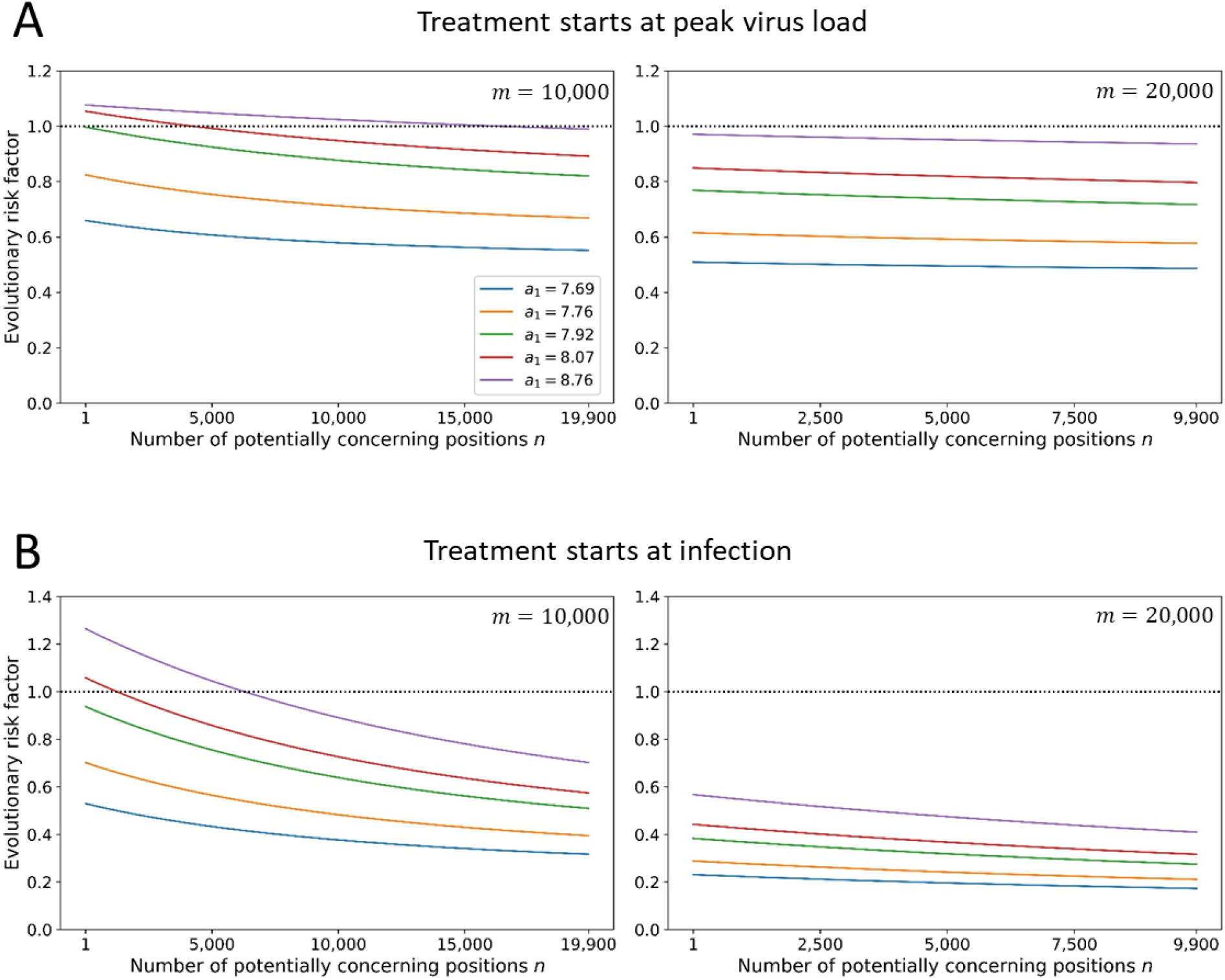
The evolutionary risk factor (ERF) versus the number of concerning mutations. The ERF of mutagenic treatment is the ratio of the cumulative mutant virus load with and without treatment. Here we show ERF versus the number *n* of potentially concerning mutations in the viral genome. We explore all values of *n* subject to the constraint that *m* + *n* remains below the length of the SARS-COV2 genome. We observe that the ERF decreases as function of *n*. (A) Treatment starts at peak virus load. (B) Treatment starts at infection. Parameters: *a*_0_ = 3, *b* = 7.61, *u*_0_ = 10^−6^, *u*_1_ = 3 · 10^−6^, *T* = 5. Initial condition: *x*_0_ = 1 and *y*_0_ = 0.

### Advantageous mutants do not substantially affect the evolutionary safety compared to neutral mutants

Concerning mutants could have an in-host advantage compared to wild type, such as faster a reproductive rate or a lower clearance rate. In **Extended Data Figure 6**, we evaluate a mutant with a 1% selective advantage in birth rate. As expected, we observe that the advantageous mutant reaches higher virus load than a neutral mutant. But we also observe: if there is a minimum increase in mutation rate that is required for evolutionary safety, then it is lower (or slightly lower) for the advantageous mutant. Therefore, a treatment that is evolutionarily safe for a neutral mutant is also evolutionarily safe for an advantageous mutant.

### Gradual activation of the immune system

So far, we have considered a sudden activation of the adaptive immune response by switching the clearance from *a*_0_ to *a*_1_ at time *T* resulting in a two-phase model of immunity. In reality, the immune response intensifies gradually over the course of the infection^43^. We explore a more gradual onset of the immune response in **Extended Data Figure 7**, where we add an intermediate phase during which the clearance rate is the arithmetic average of *a*_0_ and *a*_1_. We find that the ERF value for the three-phase immunity is very close to and bounded by the ERF values found for corresponding two phase simulations.

#### A simple approach captures the essence of mutagenic treatment and evolutionary safety

We further simplify our mathematical framework to obtain quantitative guidelines about the evolutionary safety of a mutagenic drug. We find that focusing on virus dynamics in the growth phase can be used to approximate the full infection dynamics, especially if the clearance rate is large. Note that clearance rates leading to infections which last longer than 100 days remain exceptions, and hence most individuals have a high clearance rate *a*_1_. The simplified approach is presented in the **Methods**. The agreement between the simplified and the full model is shown in **Extended Data Figure 8**.

The eventual goal of all mutagenic treatments would be to prevent the exponential expansion of the virus even before the onset of adaptive immunity. Using the SARS-COV2 estimates, *m* = 20,000, *b* = 7.61 and *a*_0_ = 3, we find that mutagenic treatment would have to achieve *u*_1_ > 4.65 · 10^−5^, which is a 50-fold increase of the natural mutation rate of the virus. If the mutagenic drug is less powerful, then it does not prevent the establishment of the infection, but it could still reduce both wild type and mutant abundance. The mutant virus load at time *T* is a one-humped function of the mutation rate with a maximum that is close to *u*^*^ = 1/(*bTm*). For *m* = 20,000, *b* = 7.61 and *T* = 5 we find *u*^*^ = 1.32 · 10^−6^. This value is close to the estimate of the natural mutation rate of the virus, *u*_0_ = 10^−6^. If *u*_0_ was greater than *u*^*^ then any increase in mutation rate would be evolutionarily safe. Otherwise, we need to calculate the condition for evolutionary safety. Let us introduce the parameter *s* with *u*_1_ = *su*_0_. The condition for evolutionary safety in the simplified model is

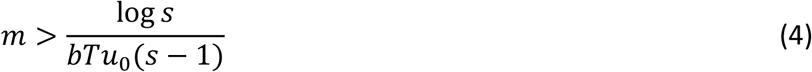

As before *b* = 7.61, *T* = 5 and *u*_0_ = 10^−6^. For *s* = 3 fold-increase of mutation rate induced by mutagenic treatment, we get *m* > 14,455. Since evolutionary safety improves with decreasing clearance rate *a*_1_ (in the full model) we can interpret inequality (4) as a sufficient condition or as an upper bound. The agreement between the analytical formulas and the numerical computation of the model is shown in **Extended Data Figure 9**. For the simplified model, we also find that ERF is a declining function of the number of concerning mutations, *n* (see **Extended Data Figure 10**).

## Discussion

We provide a mathematical framework to compute the evolutionary risk factor of death caused by mutagenic drugs and apply it to Molnupiravir, SARS-COV2 and COVID-19. For our current estimates of the parameter space, Molnupiravir treatment appears to be evolutionarily safe and can be encouraged for individuals with low clearance rates. For individuals with high clearance rates, the treatment might increase the rate of emergence of new VOCs by a few percent. However, the excess of mutant produced by immunocompetent individuals upon treatment is small in absolute amount due to the relatively smaller cumulative mutant virus load generated in such individuals.

Mutagenic treatment acts to decrease the total virus load by causing lethal mutations. It can also decrease the mutant load since (i) it eliminates the ancestors of viable mutants and (ii) it accelerates the demise of their offspring by inducing lethal mutations. In immunocompromised individuals, for which the cumulative virus load without treatment is high, mutagenic treatment can substantially reduce the amount of mutant virus generated over the course of an infection. In immunocompetent individuals, the positive effect of mutagenic treatment on reducing virus load is smaller and the abundance of mutant virus can even be increased.

Our knowledge about SARS-COV2 is still evolving. Hence, estimates for key parameters, such as the number of positions that are lethal when mutated, could change. If new estimates were to show that the value of m is below 12,000, then we predict that the evolutionary risk factor of Molnupiravir exceeds 1 and hence the treatment could increase the rate of appearance of new VOCs. We therefore advocate caution when drawing conclusions about Molnupiravir’s safety. However, our analysis has also identified parameters which will not affect appreciably the assessment of evolutionary safety of Molnupiravir, such as the number of positions that are able to give rise to new variants of concern.

Our analysis has also provide a simple rule (eq 4) for evolutionary safety of mutagenic treatment. We anticipate that additional “death by mutagenesis” drugs will emerge, and their evolutionary safety will need to be assessed before making them available for treatment.

The safety concerns that emerge from the use of a mutagenic drug extend beyond the increased rate of appearance of new VOCs. Additional deleterious effects of Molnupiravir may include the mutagenesis of the host DNA following metabolic conversion of the drug into 2′-deoxyribonucleotide^10^ and putative toxic effects on transcription of the host RNA. In addition, mutagenic treatment can have off-target effects in the event of coinfection with several pathogens. These other toxic effects are outside the scope of the current study.

Finally, the framework presented here is general enough for the assessment of evolutionary safety of this and other mutagenic drugs, in the treatment of other infectious diseases and their pathogens. Our analytical and simulation code is available on-line for further explorations (see **Code Availability**).

## Data availability

Our code for mathematical simulations is available at https://github.com/gabriela3001/molnupiravir_evol_safety. No biological data were generated during this project.

## Methods

We denote by *x* and *y* the abundances of wild type and mutant virus in an infected person. Evolutionary dynamics can be written as

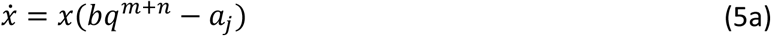

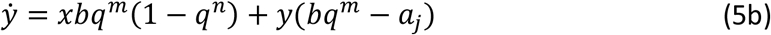

The parameter *b* denotes the birth (or replication) rate of the virus. The parameter *a*_*j*_ denotes the death (or clearance) rate of the virus. The subscript *j* indicates the absence (*j* = 0) or presence (*j* = 1) of an adaptive immune response. We have *a*_1_ > *b* > *a*_0_. The accuracy of viral replication is given by *q* = 1 − *u*, where *u* is the virus mutation rate per base. The number of lethal (or highly deleterious) positions in the viral genome is given by *m*. The number of positions in the viral genome leading to concerning mutations is given by *n*. Therefore, *y* measures the abundance of concerning mutants in a patient. At first, we assume that those mutations are neutral in the sense of having the same parameters *b* and *a*_*j*_ as the wild type virus in the patient in which they arise. We note that in Eq. (5) the mutant is mildly advantageous because *q*^*m*^ > *q*^*m*+*n*^. We assume that the adaptive immune response begins *T* days after infection, at which time the clearance rate of the virus increases from *a*_0_ to *a*_1_. Therefore, peak virus load is reached at time *T*. For exponential increase in virus load during the growth phase, which occurs during the first *T* days of infection, we require 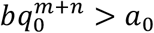. For exponential decrease in virus load during the clearance phase, we require 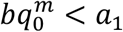.

Using *v* = *x* + *y* for the total virus abundance we obtain

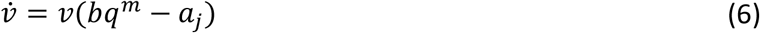

Eq (6) is the same as Eq (5a) but *m* occurs instead of *m* + *n*. In the following we derive results for *v*. The corresponding results for *x* are obtained by replacing *m* with *m* + *n*. Results for *y* are given by *v* − *x*. During the growth phase, we have 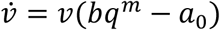. For initial condition *v* = 1 we get

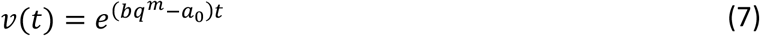

The cumulative amount of virus produced until time *T* is

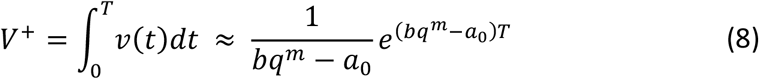

The growth phase ends at time *T*, at which point the virus abundance is

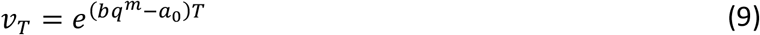

We use *v*_*T*_ and the corresponding quantities *x*_*T*_ and *y*_*T*_ as initial conditions for the clearance phase. For the clearance phase, which starts at time *T*, we have 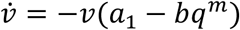. Using initial condition *v*_*T*_ we obtain

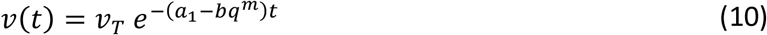

The cumulative virus during the clearance phase is given by

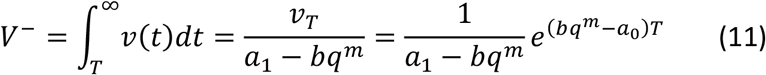

For the cumulative virus load of growth plus clearance phase we obtain

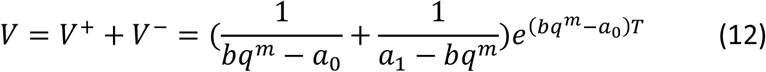

Let us use *V*_*ij*_ to denote the cumulative virus during the entire infection where *i* = 0 or *i* = 1 indicates absence or presence of treatment during the growth phase and *j* = 0 or *j* = 1 indicates absence or presence of treatment during the clearance phase. We have

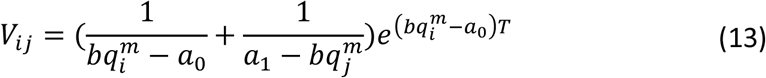

The corresponding equation for the cumulative wild type virus is

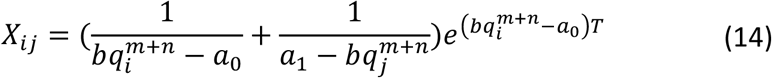

The corresponding equation for the cumulative mutant virus is given by the difference

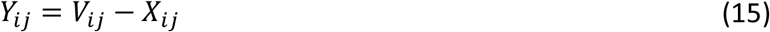

Without any treatment the cumulative mutant virus is *Y*_00_. If treatment starts at at time *T* the cumulative mutant virus is *Y*_01_. If treatment starts at time 0 the cumulative mutant virus is *Y*_11_. Mutagenic treatment increases the mutation rate of the virus from *u*_0_ to *u*_1_ and therefore reduces the replication accuracy from *q*_0_ to *q*_1_. We have *u*_0_ < *u*_1_ and *q*_0_ > *q*_1_.

### Evolutionary risk factor

We define the evolutionary risk factor, *ERF*, of mutagenic treatment as the ratio of cumulative mutant virus load with treatment over the cumulative mutant virus load without treatment. For treatment that starts at time *T*, we have *ERF* = *Y*_01_/*Y*_00_. For treatment that starts at time 0, we have *ERF* = *Y*_11_/*Y*_00_. The *ERF* quantifies how safe or unsafe a mutagenic treatment is. If *ERF* < 1 then the treatment is evolutionarily safe.

### Infectivity risk factor

We define the infectivity risk factor, *IRF*, of mutagenic treatment as the ratio of cumulative virus load with treatment over the cumulative virus load without treatment. For treatment that starts at time *T*, we have *IRF* = *V*_01_/*V*_00_. For treatment that starts at time 0, we have *IRF* = *V*_11_/*V*_00_.

### Treatment starts at peak virus load, *t* = *T*

The cumulative virus during the clearance phase with treatment is

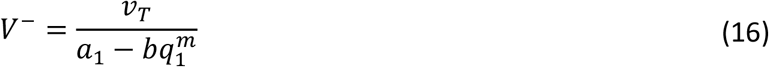

The cumulative wild type virus during the clearance phase with treatment is

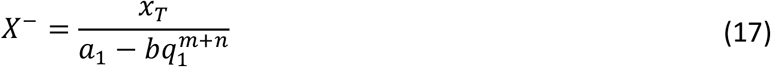

The cumulative mutant virus during the clearance phase with treatment is

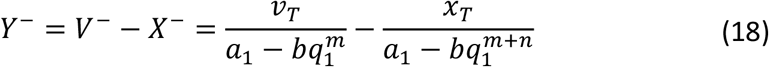

We use from above 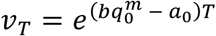 and 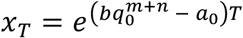. Clearly, *y*_*T*_ = *v*_*T*_ − *x*_*T*_. Let *η* = *y*_*T*_/*x*_*T*_. The function *Y*^−^(*u*_1_) has the following behavior:

1. If *η* > *n*/*m* then *Y*^−^(*u*_1_) is a declining function. In this case, mutagenic treatment is always beneficial.
2. If *η* < *n*/*m* then *Y*^−^(*u*_1_) has a single maximum which is attained at

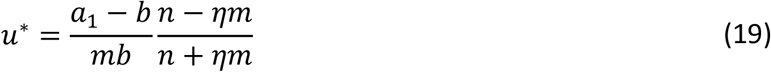

If *u*_0_ > *u*^*^ then any mutagenic treatment is beneficial. If *u*_0_ < *u*^*^ then mutagenic treatment needs to be sufficiently strong to be beneficial; in this case, we need *Y*^−^(*u*_0_) > *Y*^−^(*u*_1_). For small *u*_0_ the condition *η* > *n*/*m* is equivalent to *bT* > 1/(*mu*_0_).

### Treatment starts at infection, *t* = 0

For relevant parameters, the cumulative mutant virus load *Y*_11_(*u*_1_) – given by Eq. (15) - as a function of the mutation rate during treatment attains a single maximum at a value *u*^*^. If *u*_0_ > *u*^*^ then mutagenic treatment is always beneficial. If *u*_0_ < *u*^*^ then mutagenic treatment needs to be sufficiently strong to be beneficial; specifically, we need *Y*_11_(*u*_0_) > *Y*_11_(*u*_1_). We obtain *u*^*^ as follows. Let *μ* = *mu*. We find *μ*^*^ = *mu*^*^ as the solution of the polynomial:

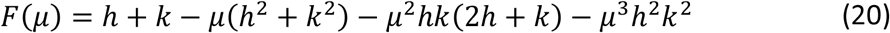

Here *h* = *bT* and *k* = [*b*(2*b* − *a*_0_ − *a*_1_)]/[(*b* − *a*_0_)(*a*_1_ − *b*)]. Exact solutions can be obtained but include complicated expressions. Approximate solutions can be found as follows. Consider fixed *h* and declining *k*. As *k* declines *μ*^*^ increases. There are 5 regions:

1. if *k* ≫ *h* then *μ*^*^ = 1/*k*
2. if *k* = *h* then *μ*^*^ = 0.52138/*k* = 0.52138/*h*
3. if *h* > *k* > 0 then *μ*^*^ < 1/*h*
4. if *h* > *k* = 0 then *μ*^*^ = 1/*h*
5. if *h* > 0 > *k* then *μ*^*^ > 1/*h* (but *μ*^*^ stays close to 1/*h*)

Therefore one can approximate as follows

1. if *k* > *h* then *μ*^*^ ≈ 1/*k*
2. if *k* > *h* then *μ*^*^ ≈ 0.52138/*h*
3. if *k* < *h* then *μ*^*^ ≈ 1/*h*

See **Extended Data Figure 1** for validity of those approximations.

### Evolutionary safety in a simplified setting

We now consider the effect of mutagenic treatment in a setting that uses further simplification. We only study the amount of virus that is generated during the growth phase with and without mutagenic treatment. As before we have:

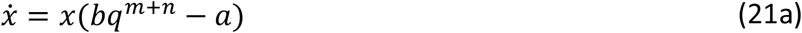

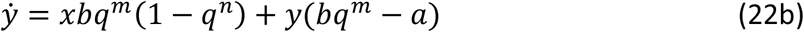

For the total virus, *v* = *x* + *y*, we have:

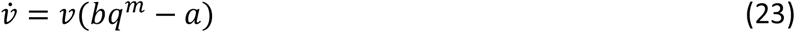

We use *q* = *q*_0_ = 1 − *u*_0_ to denote absence of treatment and *q* = *q*_1_ = 1 − *u*_1_ to denote presence of treatment, with *u*_1_ > *u*_0_. In the absence of treatment, we assume 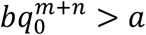 which means the wild type can expand.

Clearly, the aim of mutagenic treatment is to eradicate the infection, that is to prevent the exponential expansion. Thus, mutagenic treatment succeeds if 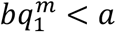. In other words, the mutation rate induced by mutagenic treatment should satisfy

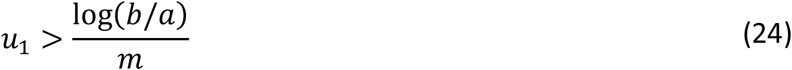

Using our SARS-Cov2 estimates, *m* = 20,000, *b* = 7.6 and *a* = 3, we obtain *u*_1_ > 4.65 · 10^−5^. If the natural mutation rate is 10^−6^ then - ideally - we are looking for a mutagenic drug that achieves a 50-fold increase in mutation rate.

If the mutagenic drug is less powerful, then it does not prevent the infection, but it could still reduce both virus load and mutant virus load. In this case a more complicated calculation is needed. For initial condition *v* = 1 (*x* = 1 and *y* = 0) we obtain at time *T*

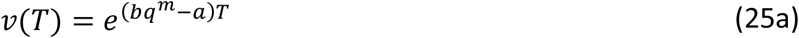

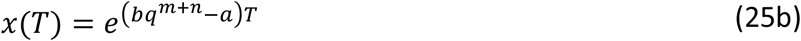

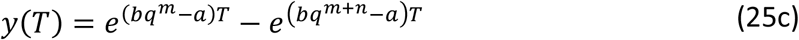

We need to understand how *y*_*T*_ behaves as a function of the mutation rate. For this analysis, the parameter *a* is irrelevant, because we can write

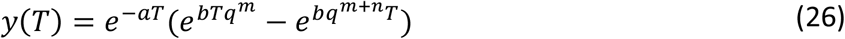

We find that *y*_*T*_(*u*) is a one-humped function with a single maximum near

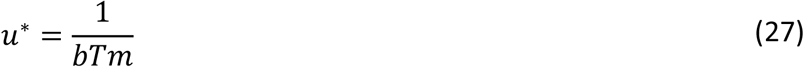

This approximation holds for *mu*^*^ << 1. Increasing *b, T*, or *m* reduces the value of *u*^*^. If *u*_0_ is greater *u*^*^ then any increase mutation rate reduces the amount of mutant virus. Using our SARS-Cov2 estimates, *m* = 20,000, *b* = 7.6 and *T* = 5, we obtain *u*^*^ = 1.31 · 10^−6^. This value is very close to the estimate for the normal mutation rate *u*_0_ = 10^−6^. If *u*_0_ is less than *u*^*^ then we need to calculate the ERF to evaluate if the treatment reduces the amount of mutant virus. We have

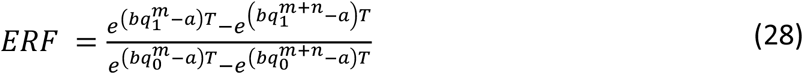

Notice that *a* cancels out and the parameters *b* and *T* appear as the product *h* = *bT*. We obtain

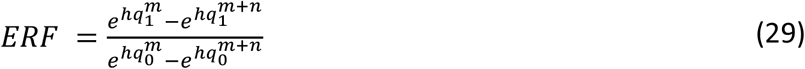

Using the approximation *q*^*m*+*n*^ = (1 − *u*)^*m*+*n*^ ≈ 1 − *u*(*m* + *n*), we get

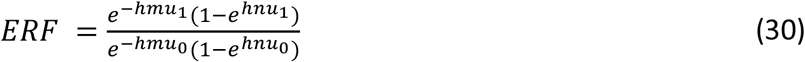

For small *hnu* we can approximate *e*^−*hnu*^ ≈ 1 − *hnu*, and therefore

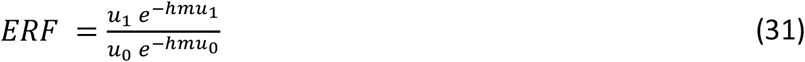

We find *ERF* < 1 if

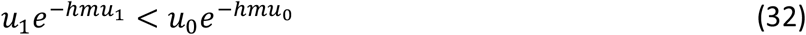

Which means

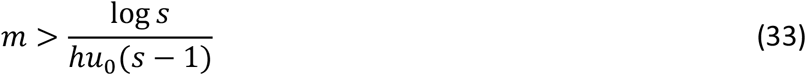

The key parameter, *h* = *bT*, is the number of replication events between the infecting virion and those virions that are present at the time of evaluation; using *b* = 7.61 and *T* = 5 we have *h* = 38.05. For *u*_0_ = 10^−6^ and *s* = 3 fold-increase induced by mutagenic treatment, we get *m* > 14,455. For *s* = 2 we get *m* > 18,217.

Defining the infectivity risk factor, *IRF*, as *v*_1_(*T*)/*v*_0_(*T*) we obtain

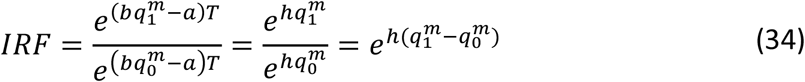

Using the approximation *q*^*m*^ = (1 − *u*)^*m*^ ≈ 1 − *mu* we have

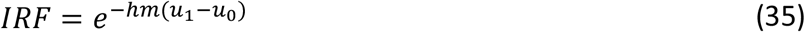

We note that IRF is always less than 1.

### Treatment increases the mutation rate only in a fraction *f* of positions

Molnupiravir is molecularly similar to a cytosine, however it can base-pair equally efficiently with both adenosine and guanosine. Hence, the probability of certain possible mutations will be increases more than others. Specifically, in the case of molnupiravir, transition mutations will be more frequent, but transversion mutations are not expected to increase. If the mutagenic drug increases the mutation rate in a fraction *f* of positions, evolutionary dynamics can be written as

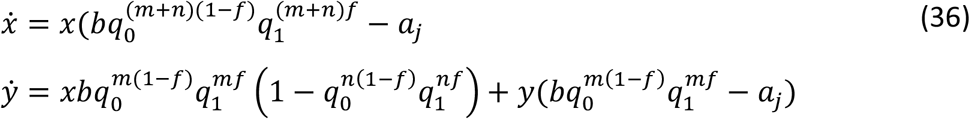

Let 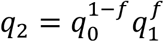. Hence, we have:

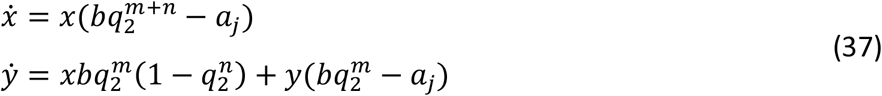

which is equivalent to Eq. 5. Hence all the subsequent derivations hold.

## Extended Data Figures

**Extended Data Figure 1.**
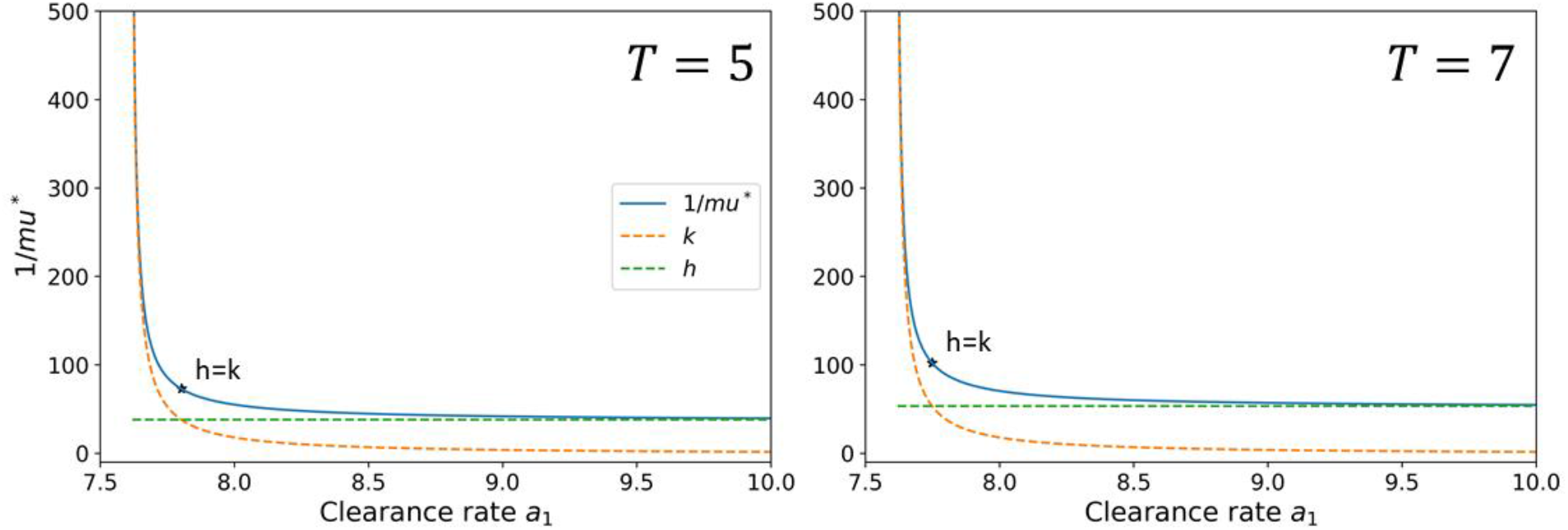
Validity of approximations for *u*^*^ when treatment starts at infection. The cumulative mutant virus load, *Y*(*u*), is a one humped function which attains a maximum at mutation rate *u*^*^. The figure shows the value of 1/*mu*^*^as function of *a*_1_. We use the notation *h* = *bT* and *k* = [*b*(2*b* − *a*_0_ − *a*_1_)]/[(*b* − *a*_0_)(*a*_1_ − *b*)]. If *h* ≪ *k* then 1/*mu*^*^ ≈ *h*. If *k* ≪ *h* then 1/*mu*^*^ ≈ *k*. If *h* ≈ *k* then 1/*mu*^*^ ≈ *h*/0.52138. We observe good agreement. Other parameters: *b* = 7.61, *a*_0_ = 3, *m* = 20,000, *n* = 1.

**Extended Data Figure 2.**
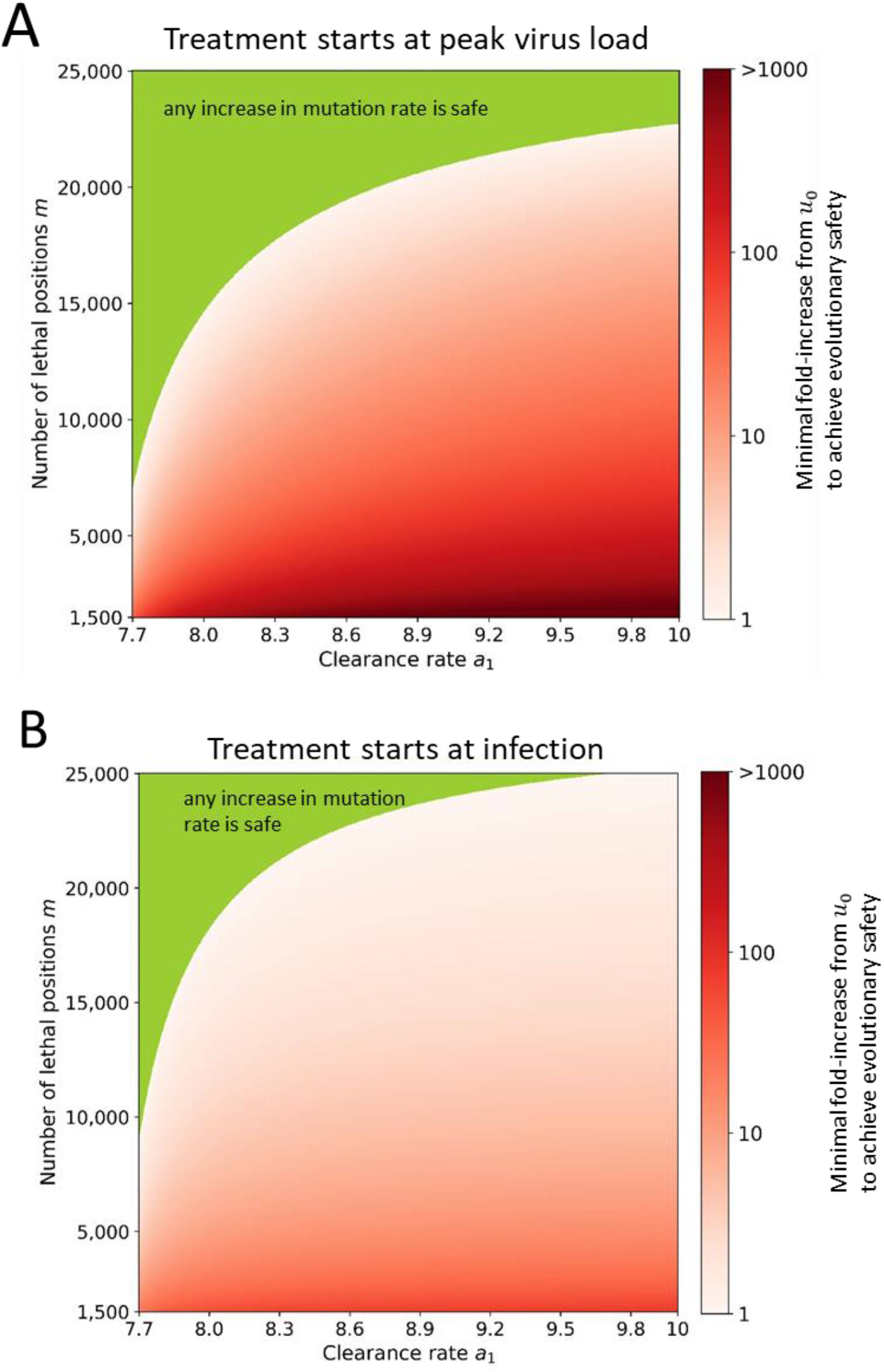
Evolutionary safety of mutagenic treatment. Same as Figure 4 of the main text but the parameter m extends to lower values. (A) Treatment starts at peak virus load. (B) Treatment starts at infection. Parameters: *b* = 7.61, *a*_0_ = 3, *n* = 1, *T* = 5, *u*_0_ = 10^−6^.

**Extended Data Figure 3.**
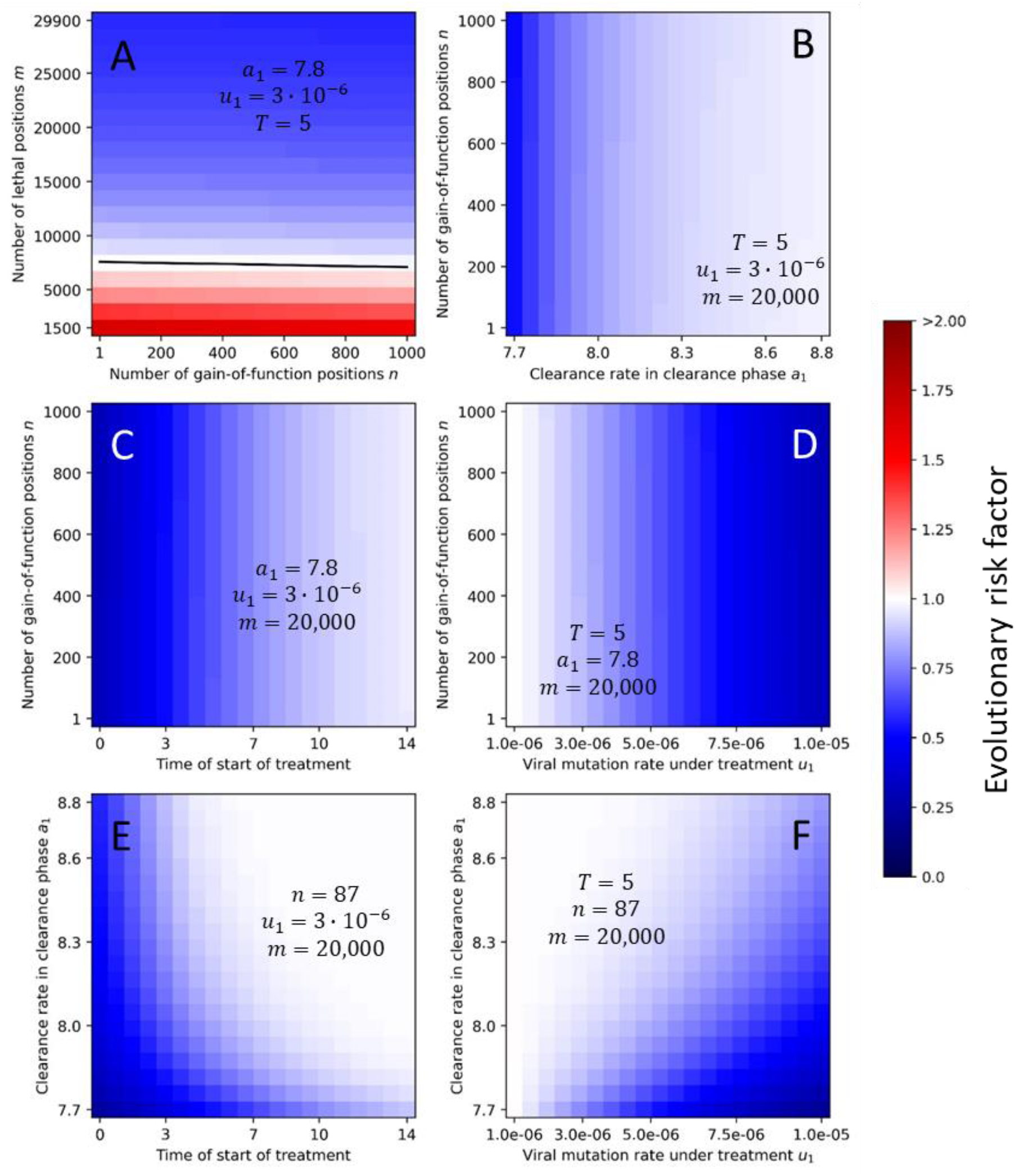
Evolutionary risk factor for a grid of pairs of selected parameters. For each pair of parameters, we numerically computed the ERF for a range of values, while other parameters were fixed. We observe that the value of *n* has little effect on the ERF. ERFs above 1 are only observed for low values of the number of lethal positions *m*. The ERF decreases with early treatment, high viral mutation rate under treatment, large number of lethal positions. Initial condition: *x*_0_ = 1 and *y*_0_ = 0.

**Extended Data Figure 4.**
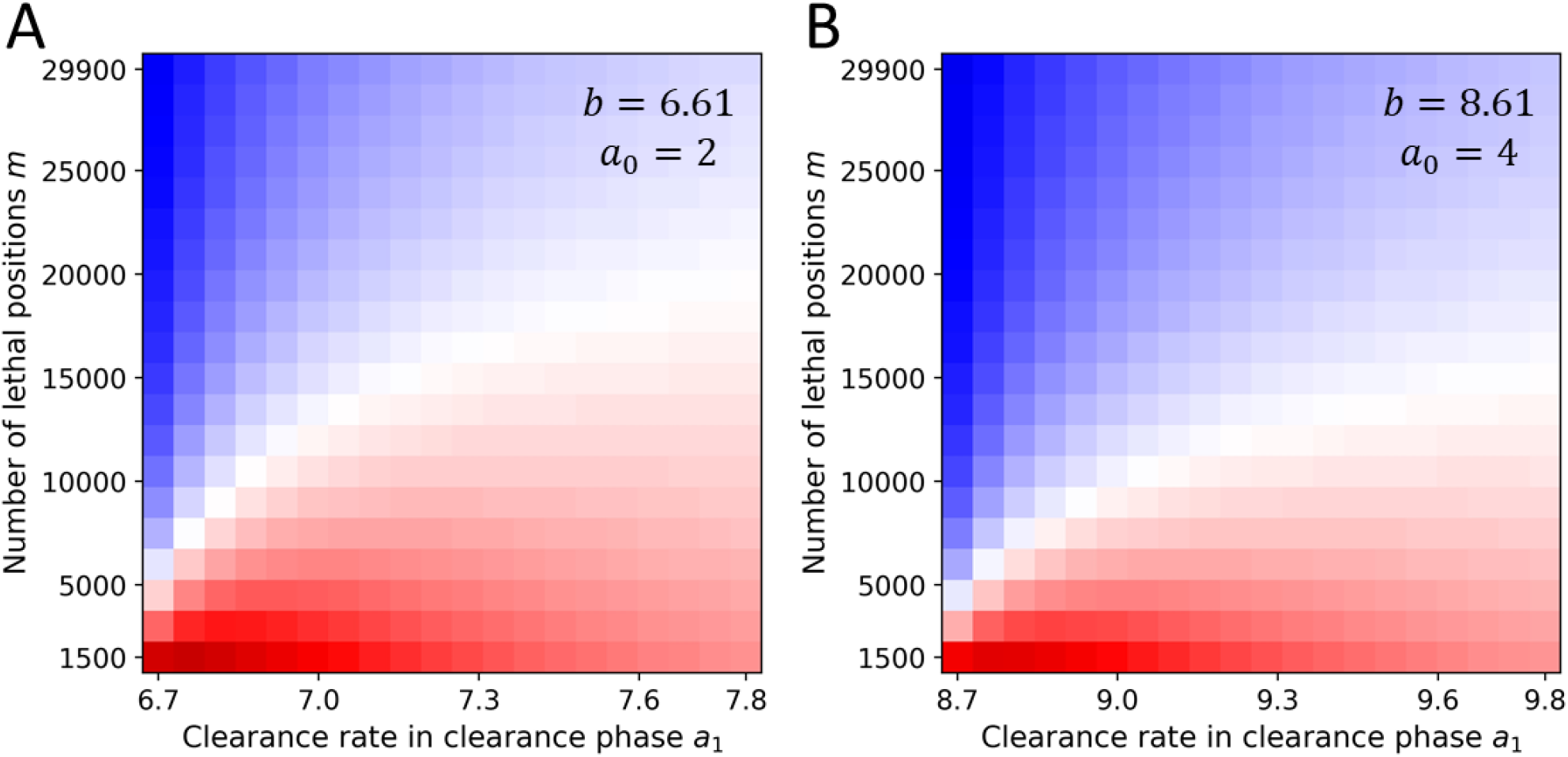
Evolutionary risk factor (ERF) for other values of *b* and *a*_0_. Our estimates for viral birth and death rates in the growth phase are *b* = 7.61 and *a*_0_ = 3. Here we compute a parameter grid of ERF versus *m* and *a*_1_for two other choices of *b* and *a*_0_, which maintain the same net growth rate (ignoring lethal mutations). For *b* = 6.61 and *a*_0_ = 2 we observe slightly higher ERF values. For *b* = 8.61 and *a*_0_ = 4 we observe slightly lower ERF values.

**Extended Data Figure 5.**
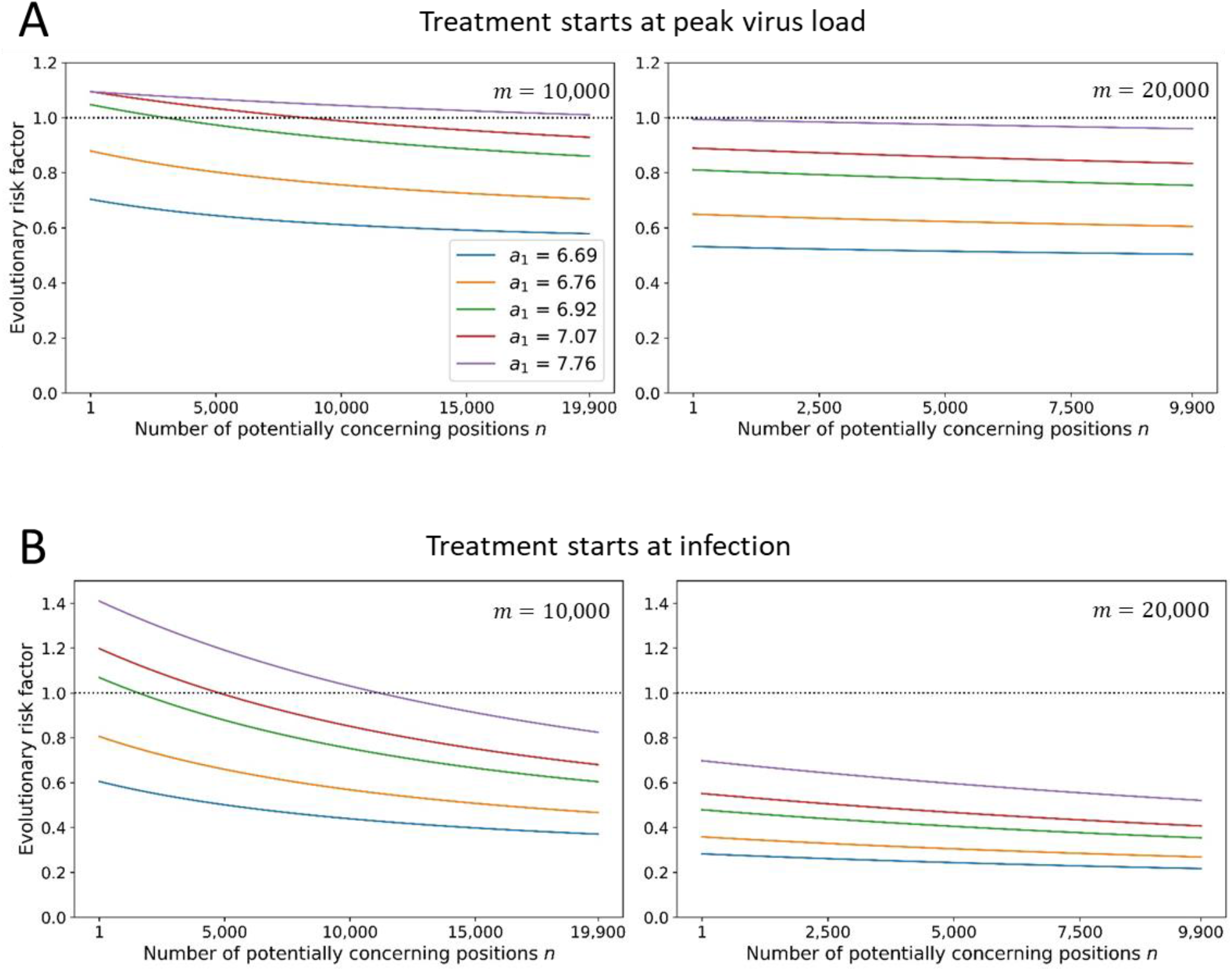
Evolutionary risk factor versus the number of concerning mutations for lower value of birth rate, *b*. Here the ERF is slightly higher than for Figure 5, which uses b = 7.61, but is still a declining function of the number of concerning positions, *n*. Parameters: *b* = 6.61, *a*_0_ = 2, *u*_0_ = 10^−6^, *u*_1_ = 3 · 10^−6^, *T* = 5. Initial condition: *x*_0_ = 1 and *y*_0_ = 0.

**Extended Data Figure 6.**
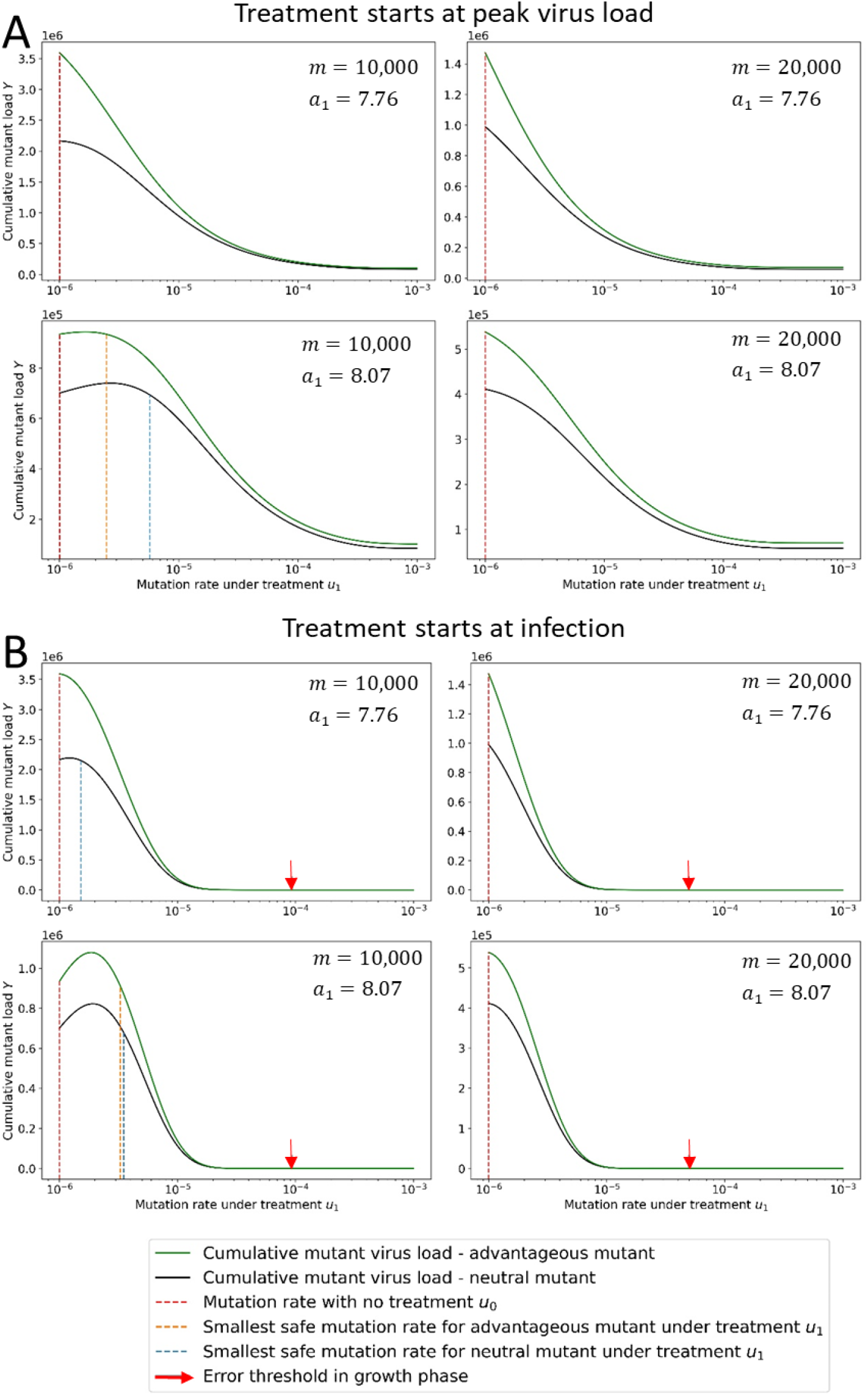
Cumulative mutant load Y versus mutation rate *u*_1_ for the case of an advantageous concerning mutant. We consider a concerning mutant with a 1% advantage in the birth rate. As expected, we observe a higher cumulative mutant load for the advantageous mutant (green line) compared to the neutral mutant (blue line). But the minimum mutation rate under treatment which is required for evolutionary safety is slightly lower for the advantageous mutant. Parameters: *b* = 7.61, *b*_*MT*_ = 7.69, *a*_0_ = 3, *n* = 1, *T* = 5, *m* and *a*_1_ as shown.

**Extended Data Figure 7.**
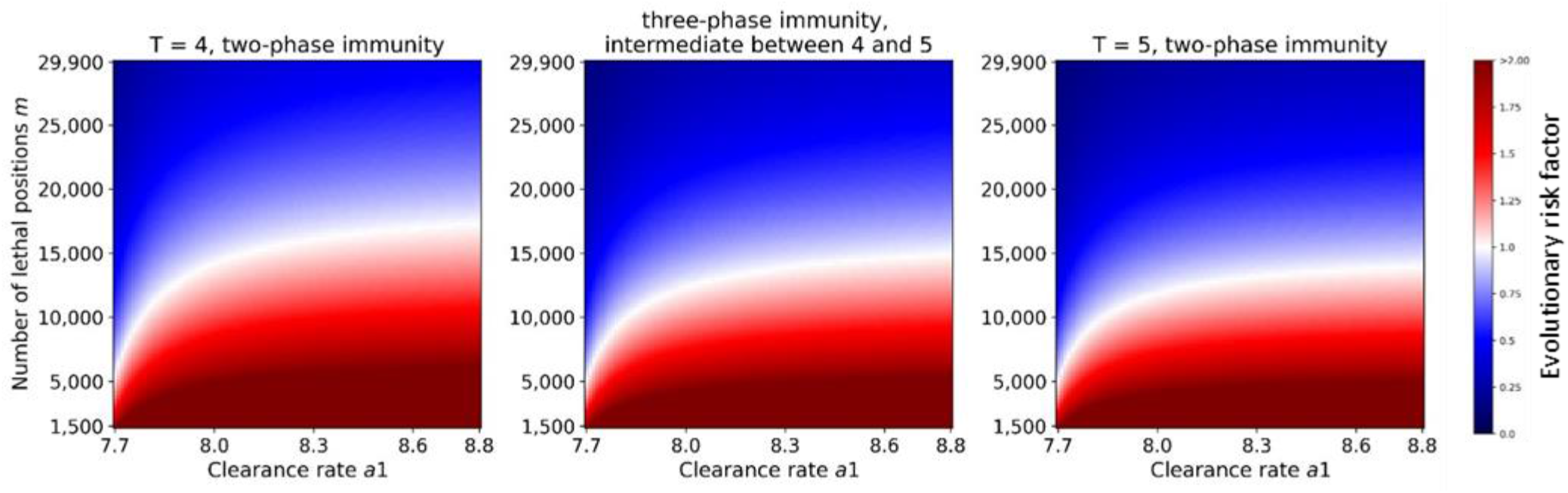
Evolutionary risk factor versus the number of lethal positions, *m*, and the clearance rate, *a*_1_, for a three-phase immune response. The values of ERF for a three-phase immunity scenario – where the clearance rate equals to the arithmetic average of *a*_0_ and *a*_1_ between days 4 and 5 – is bounded from below by the ERF values of two-phase immunity with T = 5 and is bounded from above by the ERF values of two-phase immunity with T = 4. Treatment starts at infection. Parameters: *n* = 1, *u*_1_ = 3 · 10^−6^. Initial condition: *x*_0_ = 1 and *y*_0_ = 0.

**Extended Data Figure 8.**
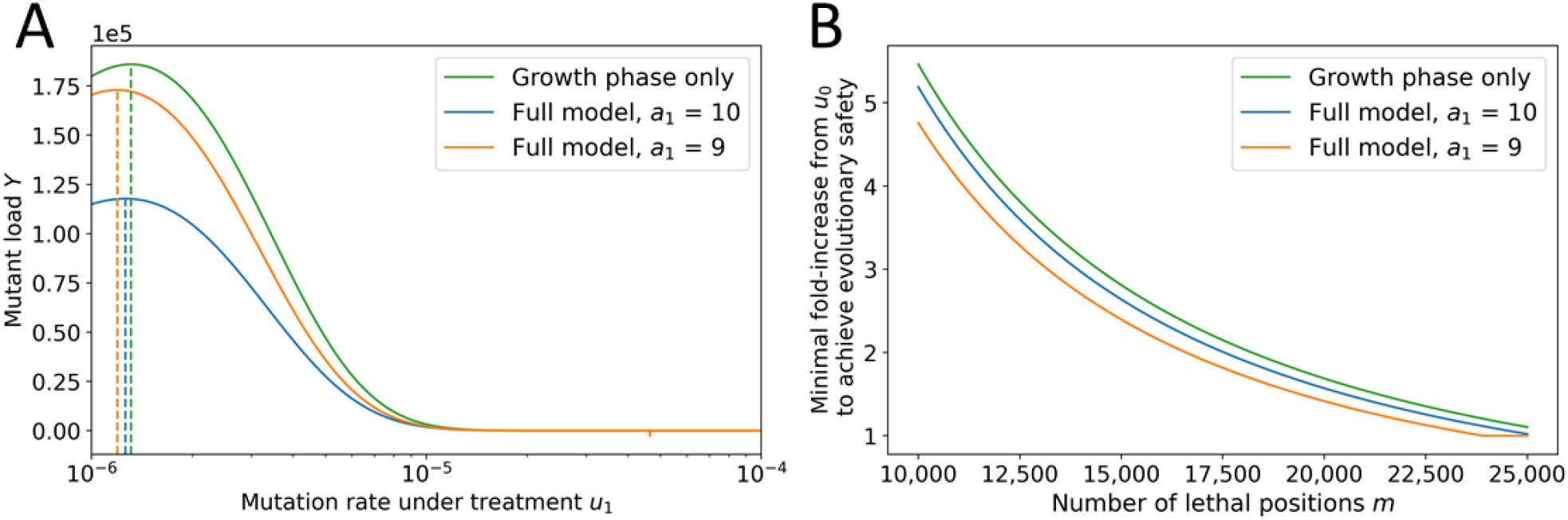
Comparing the simplified and the full model. In the simplified model, we only consider the growth phase of the virus, and we use for evaluation the abundance of mutant virus at time T. (A) Comparison between abundance of mutant virus at the end of the growth phase (green line) and the cumulative mutant virus load of the full model (orange and blue lines). The mutation rates at peak are indicated with a dashed line and are very close. (B) Minimum fold increase of mutation rate which treatment must induce to be evolutionarily safe. The simple model (green line) is a good approximation for the full model with fast clearance rates. Parameters: *b* = 7.61, *a*_0_ = 3, *n* = 1, *T* = 5, and *m* = 20,000. Initial condition: *x*_0_ = 1 and *y*_0_ = 0.

**Extended Data Figure 9.**
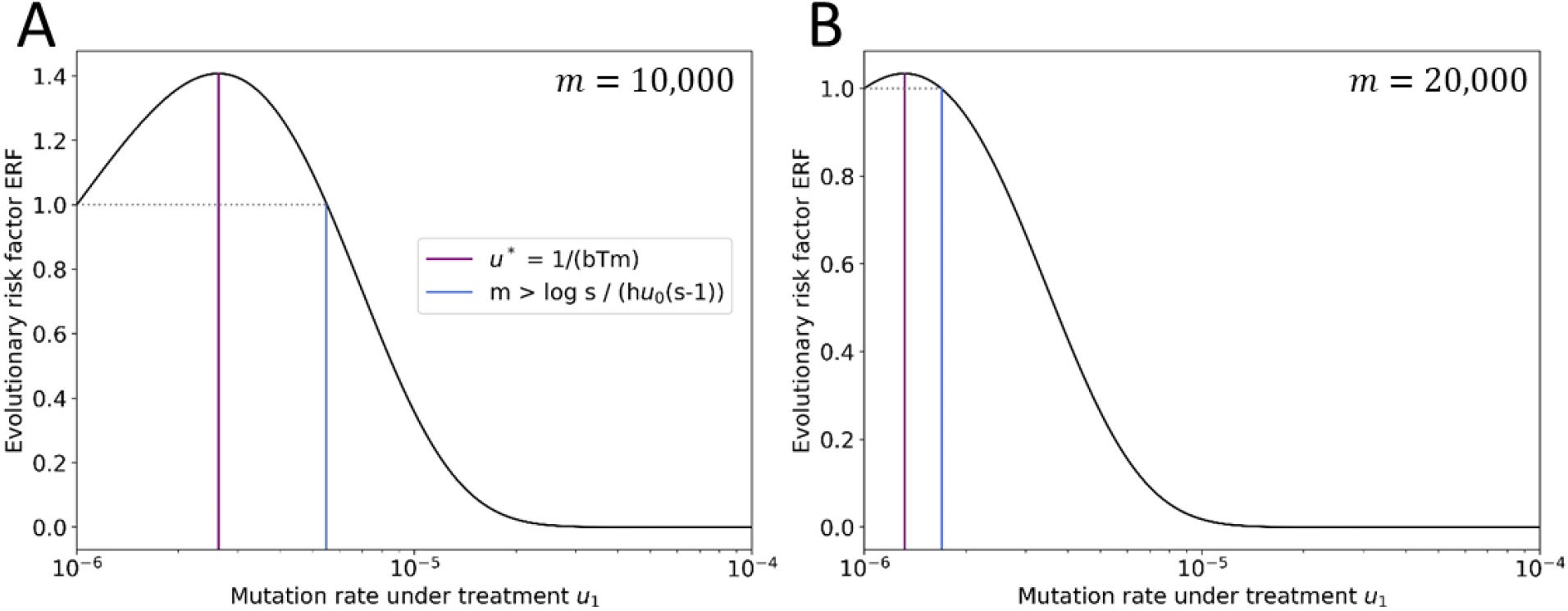
Agreement between the analytical formulas for *u*^*^ and for the minimum evolutionarily safe mutation rate under treatment considering the growth phase only. The formulas represented by the purple and blue line correspond to Eqs. 27 and Eq. 33 in the Methods. We observe perfect agreement. Parameters: *u*_0_ = 10^−6^, *u*_1_ = 3 · 10^−6^, *b* = 7.61, *a* = 3, *T* = 5 and *m* as shown.

**Extended Data Figure 10.**
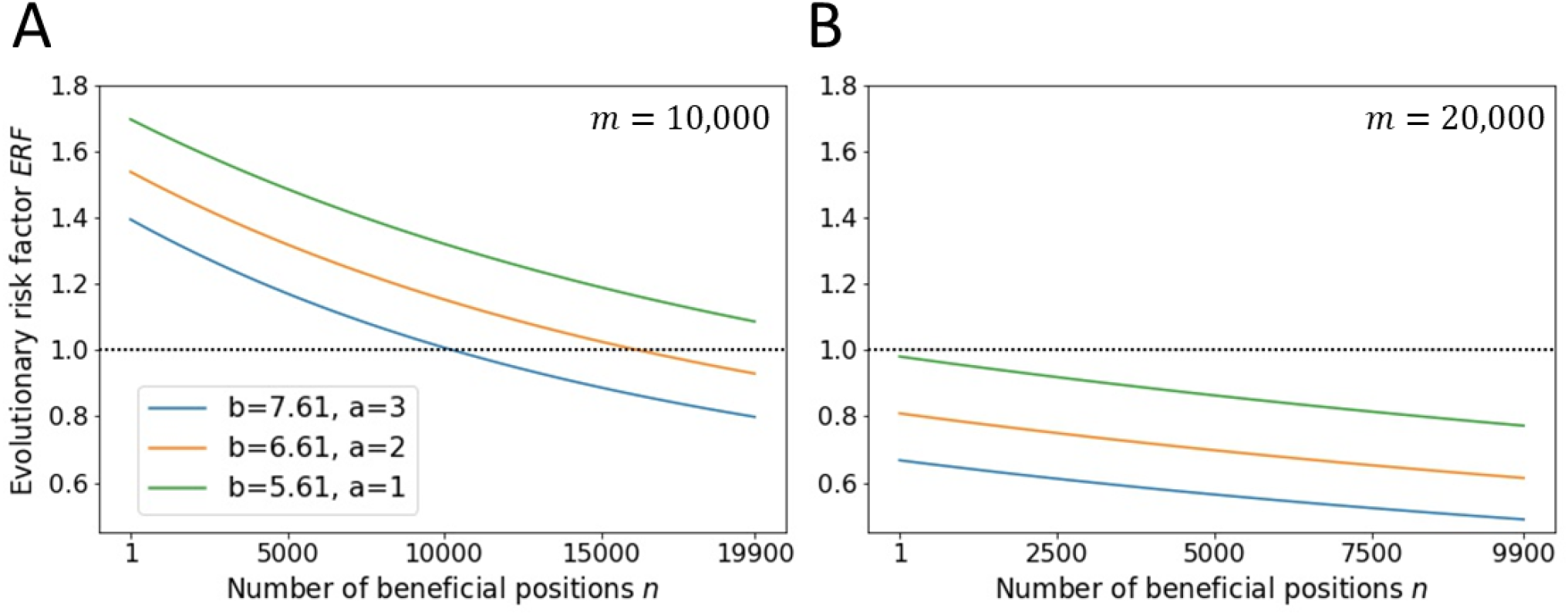
The evolutionary risk factor is a declining function of the number of concerning mutations *n* also in the simplified setting. ERF (as given by Eq. 31 in Methods) is a declining function of *n*. Parameters: *u*_0_ = 10^−6^, *u*_1_ = 3 · 10^−6^, *b, m*, and *a* as shown.

